# Micronuclear collapse under oxidative stress drives amphisome-mediated export of DNA in Parkinson’s disease

**DOI:** 10.64898/2025.11.30.691376

**Authors:** H Tavares, O Gonçalves, R Gaspar, M Beatriz, CM Deus, S Benfeito, F Cágide, F Borges, P Seibler, C Klein, AR Esteves, SM Cardoso, N Raimundo, I Milosevic, P Pinheiro, PJ Oliveira, C Lopes

## Abstract

The diversity of extracellular vesicle (EV) subpopulations and their impact on intercellular communication are increasingly recognized, but how organelle dysfunction shapes EV content in neurodegenerative diseases remains unclear. Mitochondrial and lysosomal functional defects are hallmarks of Parkinson’s disease (PD). Here we uncover a novel pathway linking this dysfunctional axis to EV remodeling and immune activation. We show that mitochondrial reactive oxygen species (ROS) induce genomic instability and micronuclei formation in PD fibroblasts, with ruptured micronuclei being sequestered into amphisomes and exported through small EVs. These EVs are enriched in oxidized mitochondrial and nuclear DNA, which potently stimulate microglial inflammatory responses. Mechanistically, this work identifies micronuclei not as passive byproducts of genome instability but as active intermediates in EV cargo loading. Importantly, treatment with the mitochondria-targeted antioxidant AntiOxCIN4 elicited a mitohormetic response, enhancing ATM-mediated DNA damage repair, restoring mitochondrial dynamics, and improving lysosomal function. This reduced the incorporation of oxidized DNA into EVs and blunted their pro-inflammatory activity. Together, our findings reveal a previously unrecognized mechanism by which mitochondrial–lysosomal dysfunction drives the release of DNA-enriched EVs that fuel neuroinflammation in a neurodegenerative context. Targeting mitochondrial quality control to limit oxidized cargo in EVs emerges as a potential strategy to mitigate early inflammatory events in PD.

## Introduction

Mitochondrial dysfunction is a hallmark of neurodegenerative diseases, including Parkinson’s disease (PD), leading to disrupted oxidative phosphorylation, excessive production of reactive oxygen species (ROS), impaired organelle dynamics, and cellular stress ^1,2^. Mitochondria have a tight functional coupling to lysosomes, organelles that beyond their traditional degradative roles also regulate cellular homeostasis, autophagy, and endolysosomal trafficking ^3^. Persistent mitochondrial ROS production, coupled with lysosomal impairment and defective mitophagy, has been linked to aging and neurodegeneration through multiple intersecting pathways ^1,4,5^.

One way these pathways may crosstalk is through extracellular vesicles (EVs). These nanoscale, membrane-bound particles are secreted by most cell types, including neurons and glia to mediate intercellular communication ^1,6^. EVs carry proteins, lipids, and nucleic acids that influence recipient cell physiology, contributing to signaling pathways ^7–9^, regulate lysosomal function, and immune responses ^10–12^. Alterations in EV biogenesis, release, or cargo composition are increasingly recognized as contributors to neurodegenerative disease pathology. Of particular relevance, EV-associated DNA (exoDNA), predominantly double-stranded and methylated resembling genomic DNA, can activate the cyclic GMP–AMP synthase (cGAS)–stimulator of interferon genes (STING) pathway in recipient cells, linking cellular stress to innate immune signaling ^13,14^.

A possible source of exoDNA are micronuclei, cytoplasmic DNA-containing structures arising from genomic instability or defective DNA repair that are increasingly recognized in neurodegenerative conditions including PD ^15–17^. Increased mitochondrial ROS can compromise micronuclear integrity, which often contains oxidized DNA, promoting rupture and release of cytosolic double-stranded DNA that engages the cGAS–STING axis ^18,19^. However, while exoDNA is acknowledged as a potent proinflammatory signal, the mechanisms determining whether genomic DNA from micronuclei is degraded in lysosomes or selectively packaged as exoDNA into EVs remain poorly understood.

Here, we address this knowledge gap by demonstrating that increased mitochondrial ROS generation in PD fibroblasts triggers DNA damage and micronuclei formation. Impaired lysosomal degradation, linked to reduced levels of the DNA-autophagy receptor LAMP2C, reroutes micronuclei-derived DNA to amphisomes, facilitating its incorporation into EVs. Because amphisomes use molecular machinery shared with EV biogenesis, we examined the DNA load, oxidative state, and microglial impact of EVs released under these conditions. Notably, mitochondrial ROS scavenging with the mitochondria-targeted antioxidant AntiOxCIN4 restored lysosomal function, reduced the incorporation of oxidized DNA into EVs, and mitigated microglial inflammatory responses. Our findings reveal a previously unrecognized mitochondrial stress–lysosome axis governing DNA packaging into EVs, with direct implications for neuroinflammation in PD.

## Results

### Mitochondrial ROS drives DNA damage and γH2AX-positive micronuclei in Parkinson’s disease

Mitochondrial-derived ROS are a major driver of genomic instability, but their contribution to micronuclei formation in PD has not been fully elucidated. To address this relationship, we analyzed multiple sporadic (sPD) and Parkin-mutant (PRKN) primary fibroblast cultures, together with age- and sex-matched controls. Fibroblasts are a robust system for investigating mitochondrial dysfunction, as they are readily accessible and faithfully reproduce the bioenergetic defects characteristic of PD neuronal cells ^20^. Under basal conditions, PD-derived fibroblasts produce more mitochondrial ROS than controls, with notably elevated hydrogen peroxide levels ranging from an 8% increase in sPD2 to a 26% increase in sPD1, a difference that was further amplified when cells were forced to rely on oxidative metabolism (**Fig. S1a, b**). Elevated ROS production correlated with increased micronuclei formation, particularly in Parkin-mutant fibroblasts (**Fig. 1a**). Although the overall size distribution of micronuclei was similar between groups, the smallest micronuclei were more frequently located at greater distances from the primary nucleus (**Fig. 1b, Fig. S2a**).

**Figure 1.**
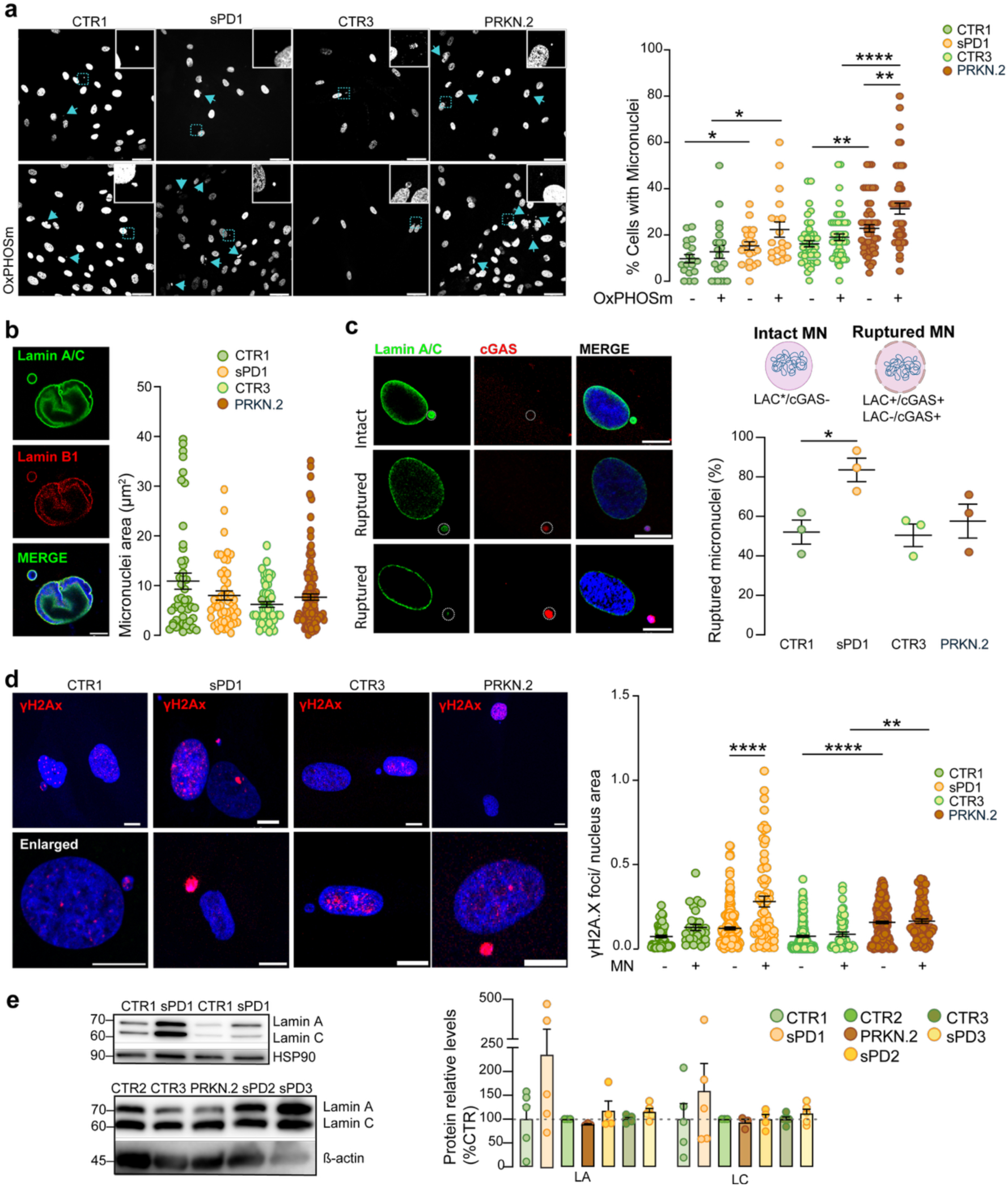
Mitochondrial ROS promote DNA damage and micronuclei formation enriched with γH2AX foci. **a,** Quantification of micronuclei formation in OXPHOS-forced sPD and PRKN fibroblasts. Arrowheads indicate representative micronuclei. Insets show enlarged images of micronuclei (n>250 nuclei; 3 independent experiments). **b,** Micronuclei area in sPD and PRKN fibroblasts. Representative image shows micronuclei stained for Lamin A/C and Lamin B1. Scale bar: 5 μm. **c,** Quantification of intact (Lamin A/C+/cGAS-) and ruptured (Lamin A/C+/cGAS+; Lamin A/C-) micronuclei in sPD and PRKN fibroblasts. Representative image shows micronuclei stained for Lamin A/C, cGAS. Scale bar: 10 μm. Data represent the percentage of micronuclei per condition (n=239 nuclei), from 3 independent experiments. **d,** Quantification of γH2AX foci in stressed fibroblasts stratified by the presence or absence of micronuclei, normalized for nucleus area. Representative image of nuclear γH2AX foci and enlarged images of γH2AX foci in micronuclei on the bottom. Scale: 10 μm. **e,** Protein levels of nuclear envelope markers Lamin A/C, HSP90 or β-actin was used as a loading control. Statistical analysis was performed using one-way ANOVA with post hoc Tukey’s test (a, d) or Mann–Whitney test (c). *p<0.05, ** p<0.01, *** p<0.001, **** p<0.0001.

To determine whether mitochondrial ROS contribute to micronuclear envelope rupture and DNA exposure at high frequency in PD cells, we labelled for Lamin A/C and cGAS in sPD and Parkin-mutant fibroblasts. Intact micronuclei retained Lamin A/C and were negative for cGAS, whereas ruptured micronuclei either lost Lamin A/C entirely or labelled positively for both Lamin A/C and cGAS, reflecting rapid recruitment of cGAS to compromised nuclear envelopes (**Fig. 1c**). We noted a significantly higher proportion of micronuclei with exposed DNA in sPD1 cells, particularly those that were positive for both Lamin A/C and cGAS, reflecting compromised micronuclei envelope integrity (**Fig. 1c; Fig. S2b**). The sPD1 cells, that exhibited higher basal ROS production, displayed a greater number of ruptured micronuclei.

Metabolic stress in PD fibroblasts led to elevated mitochondrial ROS production, which induced DNA double-strand breaks, as evidenced by increased γH2AX levels (**Fig. 1d**). This oxidative DNA damage contributes to the formation of micronuclei, a hallmark of genomic instability. Notably, a significantly higher proportion of the micronuclei observed in stressed PD cells were γH2AX-positive and contained a greater number of nuclear γH2AX foci indicating the presence of unrepaired DNA lesions. These results support a mechanistic link between mitochondrial ROS and the generation of damaged micronuclei, indicative of genomic instability.

Given the role of nuclear envelope proteins in maintaining genome integrity, we measured Lamins A, C, and B1 levels. We found elevated levels of Lamin A and C that likely represent a compensatory response to DNA damage and nuclear envelope stress; however, such increases were insufficient to restore nuclear integrity (**Fig. 1e, S2c**). These findings point to a dysregulated nuclear envelope remodeling process contributing to micronuclei persistence and defective DNA compartmentalization.

### Lysosomal impairment reroutes micronuclei into amphisomes

To investigate how micronuclei are trafficked for degradation, we assessed their engagement with the autophagic and endosomal machinery in a sporadic model of PD cells that exhibited elevated micronucleus formation, sPD1 and respective control. Using high-resolution confocal microscopy, we found that micronuclei in sPD1 fibroblasts were intimately encased by Rab7 and LAMP2, canonical markers of late endosomes and lysosomes, respectively, indicating their incorporation into the autophagic–endolysosomal pathway **(Fig. 2a).** Interestingly, we also observed that CD63, a tetraspanin associated with multivesicular endosomes (MVEs), also accumulated around micronuclei **(Fig. 2b**), suggesting the involvement of MVE-derived compartments in micronuclei degradation. Further analysis showed that micronuclei did not colocalize with markers of recycling endosomes (Rab11), secretory vesicles (Rab27), or with LAMP1 or dextran-AF647-labeled lysosomal structures, this later suggesting that they bypass canonical degradative lysosomal pathways **(Fig. S3a-f**). These findings show that micronuclei are selectively trafficked into CD63⁺/Rab7⁺/LAMP2⁺ vesicles, bypassing canonical degradative pathways. This suggests that the fusion of autophagosomes with MVEs may represent a key step in micronuclei sequestration and their release via EVs.

**Figure 2.**
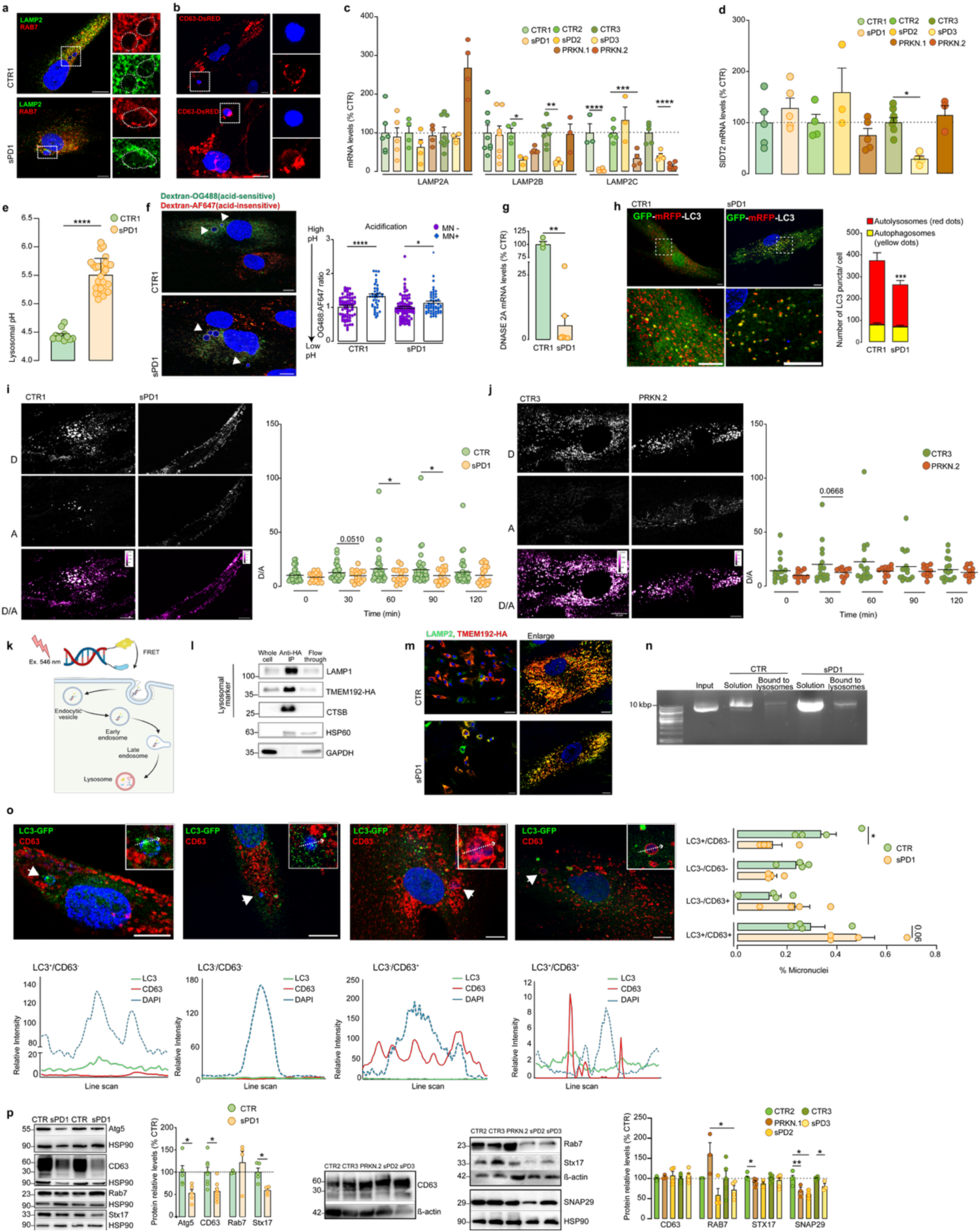
Autophagic-lysosomal dysfunction promotes micronuclei sequestration into amphisomes. **a,** Micronuclei formed in OXPHOS-forced sPD and PRKN fibroblasts colocalize with vesicles positive for LAMP2 or RAB7, indicating recruitment of endolysosomal compartments. Insets shown at right. **b**, Micronuclei also colocalized with CD63-positive vesicles, indicating their association with multivesicular bodies or late endosomal compartments. Insets shown at right. Scale: 10 μm. **c-d,** mRNA expression levels of lysosomal-associated genes LAMP2A, LAMP2B, LAMP2C, and SIDT2. **e,** Quantification of lysosomal pH using the ratiometric dye LysoSensor Yellow/Blue DND-160. Calibration curve was generated as described in the Methods. **f,** Lysosomal acidification status of sPD and control fibroblasts, stratified by the presence of micronuclei, using dextran-Oregon Green 488 (OG488) and dextran-Alexa Fluor 647 (AF647). Increased luminal pH is reflected by a higher OG488:AF647 ratio. Arrows indicate micronuclei. **g,** mRNA expression levels of lysosomal-associated gene DNASE2a. **h,** Autophagic flux assessed by mRFP-GFP-LC3. The numbers of autophagosomes (green puncta) and autolysosomes (yellow puncta) in each cell were quantified (total of 63 CTR and 66 PD from 3 independent experiments). Scale: 10 μm. **I-j**, Quantification of lysosomal donor/acceptor (D/A) ratios over time in control and PD cells (4 h pulse). Scale represents calibration of D/A values. The scale is represented by the calibration bar (D/A values). Scale: 10 μm. **k,** Schematic representation of DNA reporter’s design and trafficking for evaluating DNA-degradation ability of lysosomes. **l,** Immunoblot analysis of protein markers from different subcellular compartments in purified lysosomes, flow-through and whole-cell fractions. Lysates were obtained from cells expressing Tmem192–3xHA. **m,** Localization of Tmem192-3xHA fusion protein to lysosomes. Immunofluorescence staining with antibodies against the HA epitope tag (red) and the lysosomal marker LAMP2 (green). Scale bars: 50 μm (overview) and 10 μm (insets). **n,** Uptake and degradation of DNA by isolated lysosomes. Purified lysosomes were incubated with 1 μg of purified DNA for 10 min at 37 °C in the presence of an ATP-regenerating system and pelleted using a magnetic rack. DNA remaining in the supernatant (outside lysosomes) and DNA associated with pelleted lysosomes were analysed by agarose gel electrophoresis. Input DNA control corresponds to 1 ug DNA incubated under identical conditions but in the absence of lysosomes. **o,** Representative images and line-scan profiles showing fluorescence intensity of LC3-GFP and CD63 vesicles. Quantification of micronuclei colocalization with vesicle populations: LC3-GFP⁺/CD63⁻, LC3-GFP⁻/CD63⁻, LC3-GFP⁻/CD63⁺, and LC3-GFP⁺/CD63⁺ (total of 66 CTR and 75 PD from 3 independent experiments). Arrows indicate micronuclei. **p,** Protein levels of autophagic-endocytic markers Atg5, CD63, and autophagosome–lysosome fusion proteins Rab7, SNAP29 and Stx17. HSP90 was used as loading control. Data are mean ± SEM. Individual data points represent biological replicates. Statistical analyses were performed using multiple t-test with Holm–Sidak correction (c); Kruskal–Wallis test with Dunn’s post hoc test (d); Welch’s t test (i, j, n) or Mann–Whitney test (e, o). *p < 0.05; **p < 0.01; ***p < 0.001; ****p < 0.0001.

To dissect the molecular machinery underlying this process, we examined the expression of key autophagy-related proteins involved in autophagosome–lysosome fusion. Given the DNA content of micronuclei, we focused on LAMP2C, which has been shown to act as a receptor for cytosolic DNA in the DNAutophagy pathway, and on LAMP2A/B to assess involvement of broader lysosomal fusion mechanisms ^21^. Notably, LAMP2C mRNA levels were significantly reduced in all PD fibroblast cultures except for the sporadic cells sPD2, whereas SIDT2 expression remained unchanged, except in sPD3, where it was significantly decreased **(Fig. 2c, d).**

In lysosomes, luminal acidification is required for efficient cargo sorting along recycling and degradative pathways. Ratiometric LysoSensor measurements revealed that fibroblasts maintained a luminal pH of approximately 4.5, whereas sPD1 cells exhibited a slightly higher pH (**Fig. 2e**). We also assessed lysosomal heterogeneity within the cell by measuring the pH of individual lysosomes based on the presence of micronuclei using ratiometric fluorescence microscopy. The lysosomes of fibroblasts were loaded with the pH-sensitive Oregon green 488–dextran and the pH-insensitive dextran conjugated to Alexa Fluor 647 probe. Micronuclei-positive cells showed a significant increase in luminal pH as revealed by the increased ratio of OG488:AF647 signal intensity reflecting heterogeneity of the fibroblast population (**Fig. 2f**). Additionally, transcript levels of DNASE2A, a lysosomal DNA endonuclease that degrades DNA to oligonucleotides and nucleotides ^22^, were reduced in sPD1 fibroblasts, supporting impaired lysosomal DNA degradation (**Fig. 2g**).

Since lysosome degradative capacity seems impaired, we further measured the autophagic flux using tandem mRFP-GFP-LC3 imaging. We observed a marked decrease in red-only puncta in sPD1 cells, probably due to arrested autophagosome-to-lysosome progression (**Fig. 2h**). To confirm these results, we monitored the lysosomal DNA degradation ability using a recently developed FRET-based DNA reporter ^23^. Across all sampled time points, we saw reduced D/A values (donor signal (D)/ FRET signal (A)) in PD cells, although statistical significance was only reached at 60 and 90 minutes for sPD1 consistent with impaired lysosomal DNAutophagy (**Fig. 2i, j**). After treatment with CQ, a significant reduction in D/A ratios over time was observed (**Fig. s3h).** A time-dependent colocalization of the DNA reporter with LysoTracker showed overlap. Interestingly, control cells, displayed a higher spatial overlap of the DNA reporter with acidic compartments, which correlates with more efficient DNA degradation, whereas PD cells exhibited reduced colocalization, indicative of impaired lysosomal uptake of DNA (**Fig. S3i).**

We next analyzed whether DNA can be directly taken up by lysosomes in an ATP-dependent manner, as previously described ^24^. Lysosomes were isolated from cells expressing Transmembrane Protein 192 (TMEM192) fused to three tandem HA-epitopes ^25^. The obtained lysosomal fraction was enriched in lysosomal markers, although low levels of mitochondrial proteins were also detected **(Fig. 2l).** TMEM192 is a transmembrane protein that retains a lysosomal localization^25^, as shown by its colocalization with lysosomal marker LAMP2 (**Fig. 2m).**

To assess direct DNA uptake, lysosomes isolated from TMEM192-HA–expressing cells were incubated with plasmid DNA in the presence of an ATP-regenerating system. Following incubation, lysosomes were separated from the surrounding medium, and DNA content was analyzed. The amount of DNA remaining outside the lysosomes was higher in the sPD fibroblasts compared with the control. Moreover, DNA associated with the lysosomal fraction (including internalized DNA) in sPD1 lysosomes appeared less degraded, suggesting impaired lysosomal DNA degradation or reduced DNA autophagy activity (**Fig. 2n**).

When lysosomal function is impaired, autophagosomes are diverted from their canonical route of fusion with lysosomes and instead fuse with MVBs to form amphisomes ^26^. These hybrid organelles contain components of both autophagic and endosomal pathways (LC3⁺/CD63⁺ hybrids) ^27^. Indeed, PD fibroblasts showed increased co-localization of LC3 with CD63-positive compartments, indicative of amphisomes formation (**Fig. S4a**). When categorizing micronuclei by LC3-GFP and CD63 status (LC3⁺/CD63⁻; LC3⁻/CD63⁻; LC3⁻/CD63⁺; LC3⁺/CD63⁺), sPD micronuclei preferentially localized to CD63⁺ compartments, regardless of LC3 presence, whereas in control cells the micronuclei localized predominantly to LC3⁺/CD63⁻ vesicles (**Fig. 2o**). Finally, immunoblotting revealed a decrease in Sxt17 and SNAP29, markers of autophagosome–lysosome fusion indicating impaired autolysosomal formation, while RAB7 levels were more variable between sPD and Parkin mutated cells (**Fig. 2p, Fig. S4b**). Together, these results indicate that amphisome-mediated sequestration of micronuclei represents a key adaptive response to lysosomal dysfunction in PD fibroblasts, connecting nuclear stress to the noncanonical export of extracellular DNA.

### Amphisome-mediated sequestration of micronuclei limits cGAS activation in PD fibroblasts

cGAS is considered an autophagic receptor for the clearance of micronuclei to dampen micronuclei-mediated innate immune responses ^28^. To determine whether amphisomal capture protects micronuclei from cGAS sensing, we analyzed sPD fibroblasts that stably expressed LC3-GFP and stained for CD63 and cGAS. Micronuclei sequestered in LC3-GFP⁺/CD63⁺ compartments, indicative of amphisomes, were almost entirely devoid of cGAS signal **(Fig 3a).** Intermediate populations (LC3⁺/CD63⁻) and micronuclei lacking both markers recruited cGAS at significantly higher frequencies, pinpointing amphisomal sequestration as a critical checkpoint that restricts cytosolic DNA sensors from detecting micronuclei-derived DNA **(Fig. 3a, Fig S5, supplementary video 1-8).** We next asked whether micronuclear size influences this protective effect. Segregating micronuclei by area revealed that smaller micronuclei (<10 µm^2^) were more prone to cGAS recruitment in both sPD and control cells **(Fig. 3b)**.

**Figure 3.**
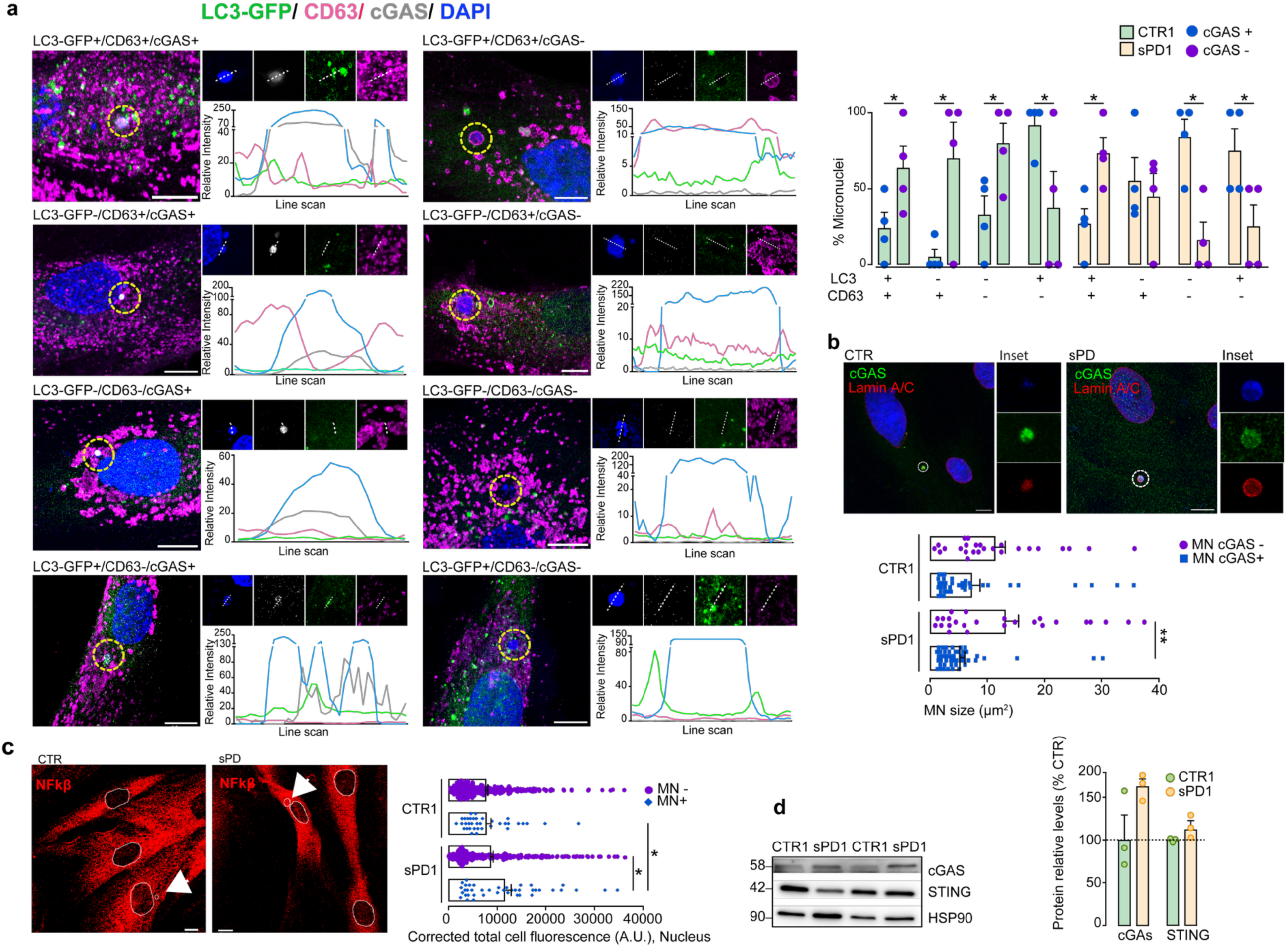
Amphisome-mediated sequestration of micronuclei limits cGAS activation in PD fibroblasts. **a,** Quantification of micronuclei categorized by LC3-GFP and CD63 status—LC3-GFP⁺/CD63⁻, LC3-GFP⁻/CD63⁻, LC3-GFP⁻/CD63⁺, and LC3-GFP⁺/CD63⁺ and their colocalization with cGAS. Representative images and line-scan profiles. Scale bar: 10 μm. The frequency of cGAS recruitment was evaluated for each subgroup. See also supplementary videos S1-8. **b**, Recruitment of cGAS in relation to micronucleus size in fibroblasts derived from sPD patients and healthy controls. Representative images show Lamin A/C⁺ micronuclei with or without cGAS signal. Scale bar: 10 μm. **c**, Nuclear translocation of NFκB in sPD and control fibroblasts, stratified by the presence of micronuclei and representative images. Scale bar: 10 μm. **d**, Immunoblot analysis of cGAS and STING protein levels in sPD and control fibroblasts. HSP90 was used as a loading control. Data are mean ± SEM. Individual data points represent biological replicates. Statistical analyses were performed using two-way ANOVA with false discovery rate (FDR) correction for multiple comparisons (a), one-way ANOVA with Tukey’s post hoc test (b) and uncorrected Fisher’s LSD test (c) *p < 0.05; **p < 0.01; ***p < 0.001.

Upon DNA binding, cGAS catalyzes the formation of cyclic GMP–AMP (cGAMP), which activates the stimulator of interferon genes (STING) located at the endoplasmic reticulum membrane. STING also activates the IκB kinase (IKK) complex, leading to the phosphorylation and degradation of IκBα, the inhibitor of NF-κB. This allows NF-κB to translocate into the nucleus and induce the expression of a broad range of pro-inflammatory cytokines, including TNF-α, IL-6, and IL-1β ^29^. To assess functional outcomes of cGAS activation, we quantified NF-κB translocation in micronuclei-bearing cells. In PD fibroblasts, cells harboring micronuclei exhibited enhanced nuclear NF-κB localization compared to micronuclei-negative cells, yet in the absence of micronuclei the overall NF-κB activity was similar between PD and control fibroblasts (**Fig. 3c**). Finally, protein analysis revealed a non-significant increase in cGAS levels in PD fibroblasts, without corresponding changes in STING levels **(Fig. 3d),** suggesting that inflammatory signaling remains modestly activated and partially buffered by amphisome-mediated sequestration. These findings establish that amphisomes may act as a selective containment system, that limits cGAS-mediated DNA sensing intracellularly, while simultaneously routing micronuclei into CD63⁺/LC3⁺ vesicles destined for extracellular release.

### Neuropathological Signatures of Autophagic and Nuclear Stress in PD Brain

To clinically validate our *in vitro* observations, we examined lysosomal and nuclear stress-related proteins in post-mortem temporal cortex samples from sporadic PD patients (Braak stage IV–VI) and age- and sex-matched controls (Table S2). Quantification of mitochondrial DNA revealed a significant reduction in mtDNA copy number in PD samples (**Fig. 4a**), in agreement with mitochondrial depletion previously reported in neurodegenerative conditions ^30^.

**Figure 4.**
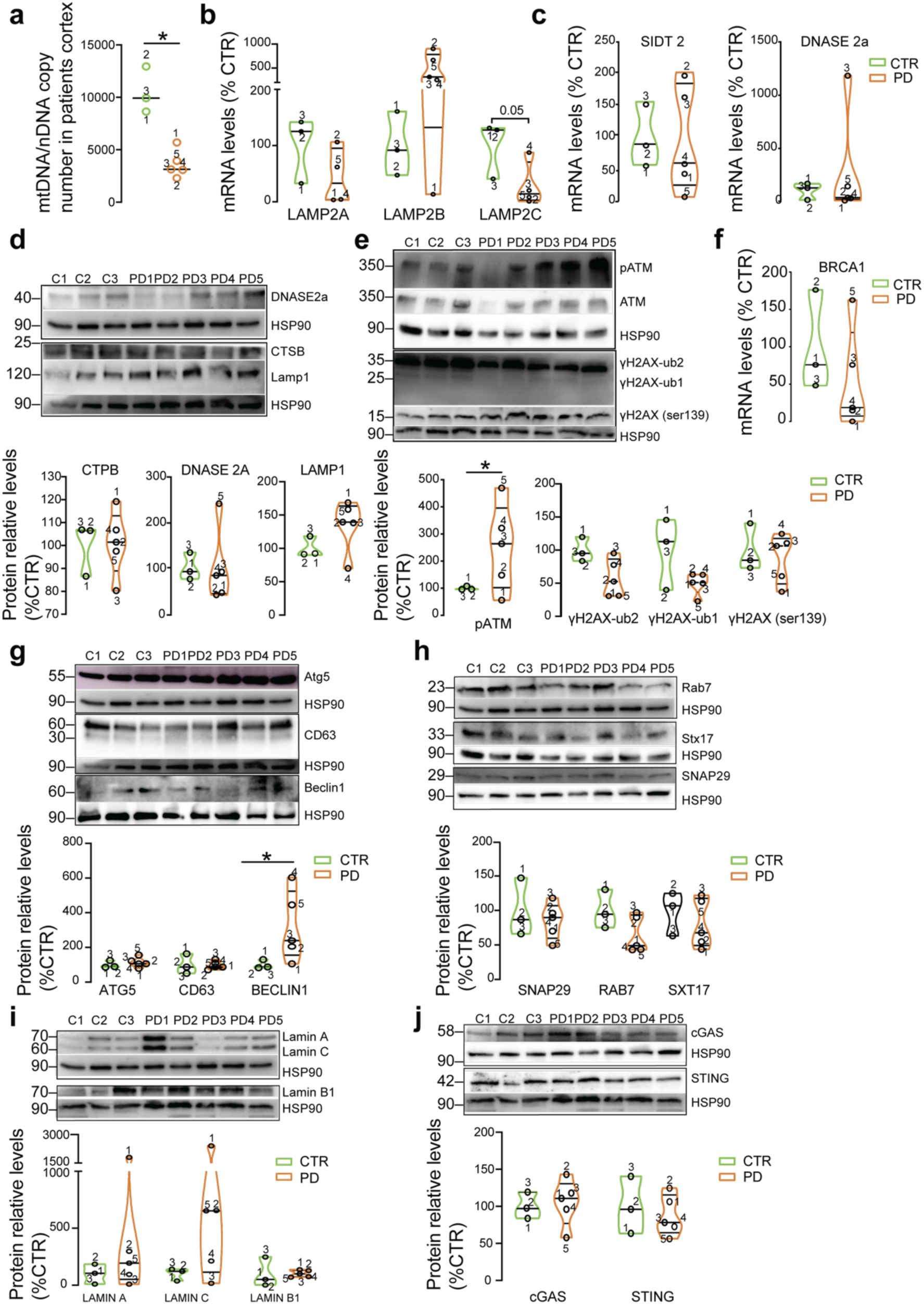
Neuropathological hallmarks in PD. **a,** Absolute mitochondrial DNA (mtDNA) copy number in post-mortem cortical brain samples from sporadic PD patients and controls. **b-c**, mRNA expression levels of LAMP2A, LAMP2B, LAMP2C, SIDT2, and DNASE2a in cortical tissue homogenates. **d,** Western blot analysis of lysosomal proteins DNASE2a, Cathepsin B and LAMP1. **e,** Western blot analysis of DNA damage response proteins: phosphorylated ATM (p-ATM), total ATM and γH2AX (mono- (ub1) and diubiquitinated (ub2) γH2AX; H2AX-Ser139-phosphorylation). **f,** mRNA levels of BRCA1. **g,** Protein levels of autophagic and endocytic markers Atg5, CD63, and Beclin-1. **h,** Levels of autophagosome–lysosome fusion proteins Rab7, Stx17 and SNAP29. **i,** Western blot analysis of nuclear envelope proteins Lamin A/C and Lamin B1. **j.** Western blot analysis of cytosolic DNA sensing proteins cGAS and STING. HSP90 was used as loading control. Each data point represents an individual patient sample. Statistical analyses were performed using Welch’s t test and the Mann–Whitney U test. *p<0.05.

We next assessed lysosomal by measuring the transcript levels of the three LAMP2 isoforms, providing insight into isoform-specific regulation. Mirroring our fibroblast data, LAMP2C mRNA was significantly reduced in sPD tissue, whereas LAMP2A showed a modest decrease and LAMP2B was elevated in nearly all patients except one (**Fig. 4b**). Other lysosomal and nucleic-acid autophagy components, SIDT2, DNASE 2a, cathepsin B (CTSB), and LAMP1, were unchanged between sPD and control samples (**Fig. 4c–d**).

To evaluate the DNA damage response, we probed ATM signaling and its downstream target γH2AX. Several sPD cases exhibited elevated phospho-ATM and increased γH2AX immunoreactivity, indicating activation of DNA damage signaling **(Fig. 4e).** However, despite this increase, the ubiquitinated form of H2A was markedly reduced, suggesting a potential defect in recruiting downstream DNA repair effectors. Supporting this, BRCA1 levels, a E3 ubiquitin ligase that targets H2A for ubiquitination and homologous recombination repair, was also significantly decreased in sPD brains. This points to a functional uncoupling between DNA damage sensing and repair execution in PD. Additionally, Beclin 1 levels were elevated, consistent with increased autophagy initiation; however, key components of the autophagosome–lysosome fusion machinery-SNAP29, STX17, and Rab7-tended to be reduced in some patients, reflecting defects in the final steps of autophagic flux. Together, these data suggest that PD brain pathology involves selective disruption of LAMP2C-dependent lysosomal degradation, impaired BRCA1-mediated DNA repair, and compromised autophagic-lysosomal fusion, linking mitochondrial and nuclear stress to impaired intracellular clearance and potential extracellular release of damaged DNA.

### Mitochondrial and lysosomal dysfunction drive DNA packaging into small EVs

To investigate how a compromised mitochondrial-lysosomal axis influences EV dynamics, we exposed normal human dermal fibroblasts (NHDF) to the mitochondrial uncoupler FCCP in combination with chloroquine (CQ), a lysosomal alkalinizing agent. This dual treatment led to mild but significant alterations in mitochondrial structure, including increased fragmentation (**Fig. S6a)**, elevated mitochondrial-derived ROS (specifically superoxide and hydrogen peroxide) (**Fig. S6b-c)** and lower mtDNA copy number (**Fig. S6d-e**), an effect further exacerbated in cells with lysosomal degradation defects

We next assessed the impact of this acute stress on small EV secretion. Small EVs were isolated from conditioned medium following 24-hour FCCP/CQ treatment using differential ultracentrifugation and characterized according to MISEV guidelines (**Fig. 5a**). Nanoparticle tracking analysis confirmed particle sizes within the expected range (**Fig. 5b**) and revealed a significant increase in small EV release upon treatment (**Fig. 5d**). Immunoblotting confirmed small EV purity, with enrichment of the canonical small EV markers flotillin-2 and CD63 and absence of the endoplasmic reticulum protein calnexin (**Fig. 5c**). Notably, despite the marked reduction in cellular mtDNA copy number, small EVs from treated cells exhibited higher mtDNA content (**Fig. 5e, f**), suggesting selective packaging or enrichment of mitochondrial components into small EVs under stress conditions. Given the oxidative stress induced by FCCP/CQ treatment, we hypothesized that this may promote oxidation of both nuclear and mitochondrial DNA. Quantification of 8-hydroxy-2′-deoxyguanosine (8-OHdG) in DNA from nuclear and mitochondrial fractions, as well as in small EV-associated DNA, revealed that mitochondrial and vesicle-associated DNA were more extensively oxidized than nuclear DNA in the treated cells (**Fig. 5g**).

**Figure 5.**
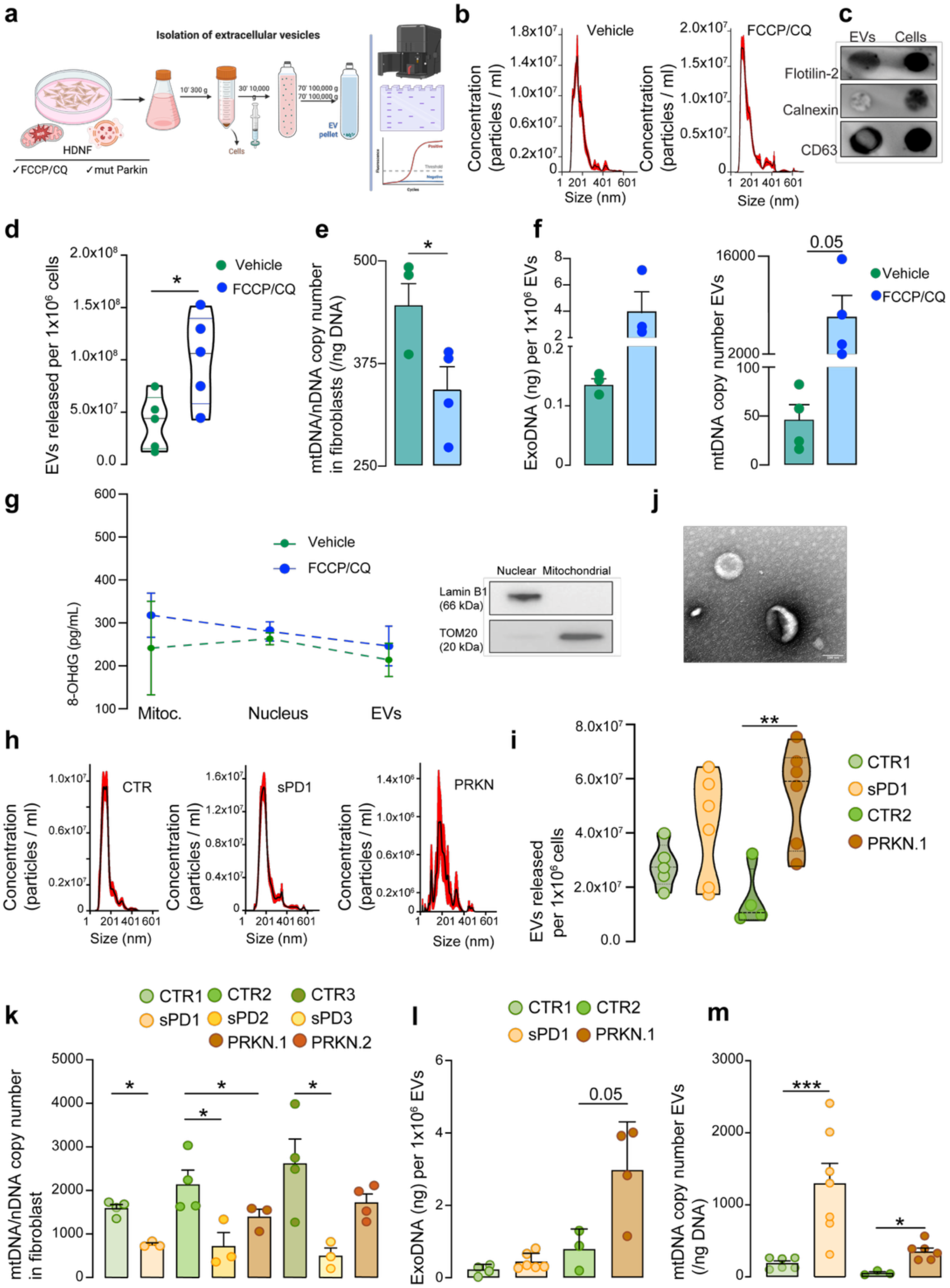
Mitochondrial-lysosomal axis dysfunction leads to increased release of sEV with oxidized mtDNA. **a,** Schematic representation of sEV isolation method and further sample processing. **b-g,** NHDF were treated with FCCP/CQ or vehicle (DMSO). **b,** NTA measurements of particle size and concentration. **c,** Dot blot analysis of fibroblast-derived sEVs showing protein markers flotillin-2 and CD63; calnexin was used as a negative control. **d,** Number of particles secreted per 1×10^6 cells. **e,** Absolute mtDNA copy number from treated fibroblasts **f**, total DNA and mtDNA copy number in corresponding purified sEV. **g,** Levels of 8-OHdG measured in mitochondrial, nuclear and EV-contained DNA samples from each condition. Representative western blot of subcellular fractionation of the mitochondrial (TOM20) and nuclear fraction (Lamin B1) of fibroblasts. **h-m,** OXPHOS-forced sPD and PRKN fibroblasts. **h**, NTA measurements of particle size and concentration. **i,** Number of particles secreted per 1×10^6 cells. **j,** TEM image of EVs. Scale: 100 μm. **k-m**, Absolute mtDNA copy number in cells and total DNA and mtDNA copy number in the corresponding purified sEVs. Bar plots represent mean ± SEM. Individual data points indicate biological replicates. Statistical analyses were performed using Student’s *t*-test (d, e, f, i) or Mann–Whitney test (k, l, m) *p < 0.05; ***p < 0.001.

Consistent with the pharmacological model, fibroblasts from sPD and Parkin-mutant patients exhibited increased basal mitochondrial ROS and impaired mitophagy under forced OXPHOS metabolism (**Fig. S1a, b**). To determine whether these pathological features similarly affect small EVs biology, we isolated and analyzed small EVs from PD fibroblasts. Consistent with the observations in treated fibroblasts, PD fibroblasts secreted higher levels of small EVs (**Fig. 5h, i**), also enriched in DNA, particularly mtDNA (**Fig. 5l, m**), despite lower cellular mtDNA levels (**Fig. 5k**). These data indicate that disrupted mitochondrial and lysosomal function drive the secretion of small EVs enriched in mitochondrial and nuclear components, including oxidized DNA. This observation aligns with our findings showing that micronuclei, generated by ROS-induced DNA damage, are rerouted into LC3⁺/CD63⁺ amphisomes rather than canonical lysosomes. The amphisome-mediated sequestration provides a noncanonical trafficking route that preserves the DNA cargo and facilitates its incorporation into small EVs.

### Mitochondria–lysosome–defective cells release bioactive small EVs that trigger microglial cytokine production

To determine whether oxidized DNA in small EVs acts as a damage-associated molecular pattern (DAMP), we treated a microglia line with one of four stimuli derived from control or FCCP/CQ–stressed fibroblasts: (i) intact small EVs, (ii) small EV-associated DNA (ExoDNA), (iii) purified mtDNA, or (iv) isolated mitochondria. After 24 hours, we quantified the mRNA levels of key proinflammatory cytokines. Microglia exposed to stimuli from FCCP/CQ–treated fibroblasts showed significantly elevated proinflammatory cytokines (TNF-α, IL-1β, IL-6) compared with untreated controls or treated with vehicle (DMSO) (**Fig. 6a–c; Fig. S7a**). Both ExoDNA and mtDNA, regardless of oxidative state, strongly induced TNF-α, whereas intact mitochondria and small EVs only triggered IL-1β and TNF-α when derived from FCCP/CQ–treated cells (**Fig. 6a-c**), highlighting the importance of cargo oxidation in modulating EV bioactivity.

**Figure 6.**
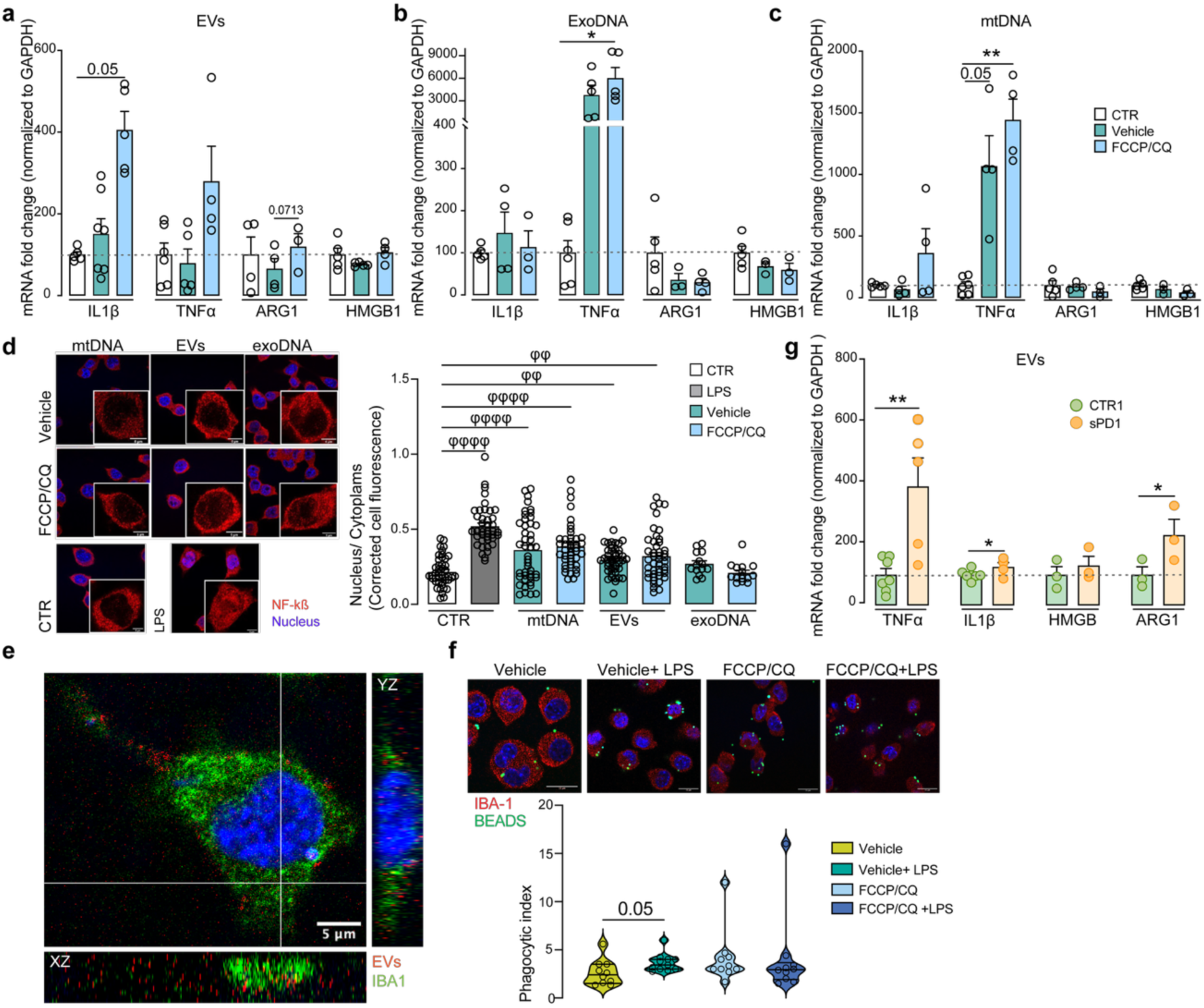
sEVs from cells with a defective mitochondria–lysosomal axis trigger pro-inflammatory cytokine release in microglia. Microglia were treated with **a,** small extracellular vesicles (sEVs) isolated from EV-depleted media of fibroblasts following FCCP or DMSO (vehicle) exposure. **b,** DNA purified from sEVs. **c,** mitochondrial DNA (mtDNA) isolated directly from fibroblasts. **d,** Nuclear translocation of NF-κB p65 in microglia treated with LPS (1 µg ml⁻¹), mtDNA, sEVs, or sEV-derived DNA. Representative confocal images show nuclear and cytoplasmic localization of NF-κB. Scale bar: 5 μm. Quantification uses 45 cells from 3 independent experiments (mean ± SEM). **e,** Confocal orthogonal projections of IBA1-positive microglia (green) internalizing mCherry-labelled sEVs (red). Scale: 5 μm. **f,** Phagocytic index (mean number of beads per phagocytic cell) in microglia treated with LPS (1 µg ml⁻¹). Scale: 10 μm. **g,** Microglia were exposed for 24 h to sEVs from Parkinson’s disease (PD) fibroblasts cultured in galactose-based, EV-depleted media. Expression (fold-change) of IL1B, TNF, ARG1 and HMGB1 mRNA, normalized to GAPDH. Data are presented as mean ± SEM (n = 3–6 independent experiments), expressed relative to control. Individual points represent biological replicates. Statistical analysis was performed using linear mixed-effects model with Tukey’s post hoc test (*p < 0.05; **p < 0.01), and Kruskal–Wallis test with Dunn’s multiple comparisons test (*φφ* p < 0.01; *φφφφ* p < 0.0001.

We then asked whether small EVs from sPD fibroblasts similarly activate microglia. Consistent with our pharmacological model, small EVs from sPD fibroblasts also elicited a robust increase in TNF-α and IL-1β expression (**Fig. 6g**). To confirm engagement of the canonical NF-κB pathway, we measured nuclear translocation of NF-κB p65. All conditions enhanced NF-κB nuclear localization, with the greatest effect observed following treatment with oxidized mtDNA and small EVs (**Fig. 6d; Fig. S7b–c**). Importantly, microglial uptake of DsRed-CD63–labeled small EVs was unaffected by cargo oxidation (**Fig. 6e–f**), indicating that heightened inflammatory responses are due to altered EV content rather than increased internalization.

Together, these findings indicate that mitochondrial and lysosomal dysfunction in donor fibroblasts not only disrupts cellular homeostasis but also reshapes the molecular composition of secreted small EVs, selectively enriching them in oxidized DNA species that function as potent DAMPs capable of activating innate immune responses in recipient microglia.

### Mitochondria-targeted antioxidant AntiOxCIN 4 reprograms EV cargo and attenuates inflammation

Given that small EVs from sPD fibroblasts are enriched in oxidized DNA and trigger microglial activation, we next explored whether targeted antioxidant therapy could reverse this pro-inflammatory small EV signature. We employed AntiOxCIN4, a mitochondriotropic compound previously shown to accumulate in the mitochondrial matrix and activate Nrf2-dependent antioxidant defenses ^31^. Beyond improving mitochondrial function, AntiOxCIN4 stabilizes redox signaling and supports lysosomal activity, two key processes disrupted in PD ^32^. Treatment of control and PD fibroblasts with AntiOxCIN4 induced a transient elevation in mitochondrial ROS, particularly hydrogen peroxide, followed by a time-dependent reduction after 48 hours, especially in control cells (**Fig. 7a, b**), as demonstrated previously^33^. PD fibroblasts showed a more attenuated decline in mitochondrial H₂O₂, consistent with persistent oxidative stress. To evaluate whether this redox modulation alters the immunogenicity of secreted vesicles, we exposed microglia to small EVs isolated from treated fibroblasts. Small EVs from AntiOxCIN4-treated PD fibroblasts, but not controls, led to a significant reduction in TNF-α and IL-1β transcript levels (**Fig. 7c**), suggesting a reprogramming of vesicle content toward a less inflammatory profile. Interestingly, AntiOxCIN4 did not significantly affect small EV secretion (**Fig. 7d**), but markedly reduced the amount of mtDNA packaged within PD-derived small EVs, despite increasing overall cellular mtDNA content (**Fig. 7e**). This suggests improved mtDNA integrity and/or containment, potentially reflecting reduced fragmentation or enhanced organellar turnover. Consistent with this, AntiOxCIN4 significantly lowered both nuclear and mitochondrial 8-OHdG levels in sPD cells (**Fig. 7f**) but not controls, indicating effective mitigation of oxidative DNA damage in PD cells.

**Figure 7.**
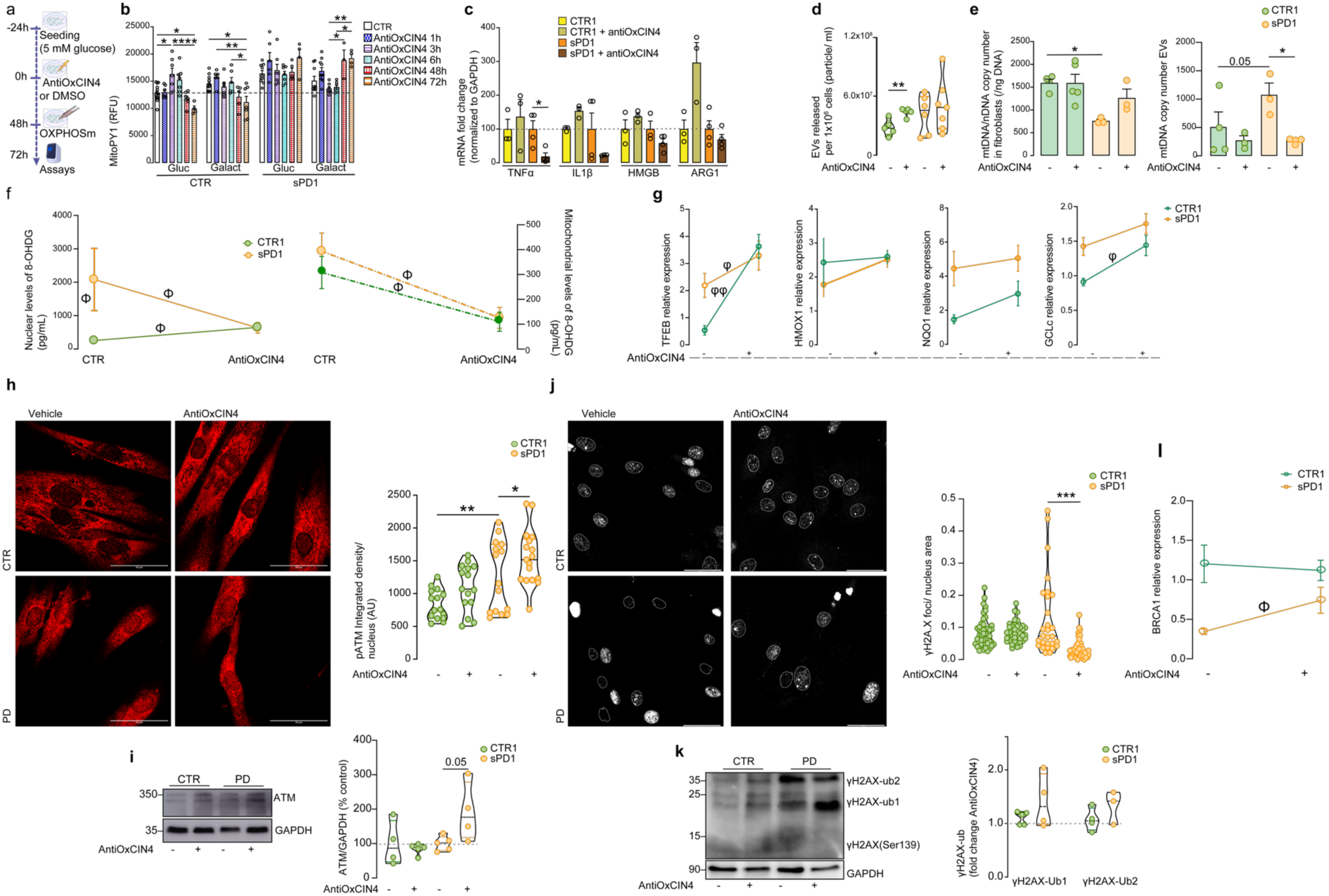
Mitochondriotropic antioxidant AntiOxCIN4 restores mitochondrial function and enhances the DNA damage response in PD fibroblasts. **a,** Schematic representation of the experimental design. **b,** Mitochondrial hydrogen peroxide production in MitoPY1-labelled cells treated with AntiOxCIN4 at different timepoints. **c,** mRNA expression of inflammatory markers (TNF-α, IL-1β, HMGB1, and ARG1) in microglia treated for 24 h with sEVs isolated from PD fibroblasts cultured in OxPHOS media for 24h. Expression levels were normalized to GAPDH and expressed relative to control. **d,** Quantification of sEV released by 1×10^6 cells. **e,** mtDNA copy number in fibroblasts (left) and their corresponding purified sEV (right). **f,** Levels of oxidative DNA damage marker 8-OHdG in the nucleus and mitochondria after AntiOxCIN4 treatment (n = 3–5). **g,** Expression of TFEB and antioxidant genes following AntiOxCIN4 treatment. **h,** Representative images of phosphorylated ATM (p-ATM) and quantification of its nuclear localization. Scale: 50 μm. **i,** Western blot analysis of total ATM levels, normalized to GAPDH. **j,** Formation of γH2AX foci: representative images and quantification. Scale: 50 μm. **k,** Detection of mono-(ub1) and diubiquitinated (ub2) γH2AX by western blot. **l,** BRCA1 mRNA-expression, normalized to GAPDH (n=3-5). Data are mean ± SEM; individual points represent biological replicates. Individual data points represent biological replicates. Statistical analysis was performed using one-way ANOVA followed by Tukey’s post hoc test (*p < 0.05, **p < 0.01, ***p < 0.001), the Mann–Whitney U test (Φ < 0.05) and Welch’ t test (***p<0.001). Panel a was created with BioRender.com.

Given the role of NRF2 in lysosomal and DNA repair signaling ^34,35^ we also assessed the transcriptional response of antioxidant and lysosomal genes. TFEB, a master regulator of lysosomal biogenesis, was significantly upregulated in both PD and control fibroblasts, whereas canonical NRF2 targets (HMOX1, NQO1, GCLc) showed modest induction (**Fig. 7g**). Notably, the AntiOxCIN4-induced upregulation of TFEB did not correlate with increased lysosomal biogenesis (**Fig. S8a),** though we did detect elevated cathepsin activity, especially in control cells, using the Magic Red cathepsin B activity assay and the acidotropic dye LysoTracker (**Fig. S8c**). Moreover, GPN-triggered calcium release confirmed improved lysosomal competence in AntiOxCIN4-treated fibroblasts (**Fig. S8b**).

Mitochondrial ROS is also a potent activator of the DNA damage response via ATM kinase ^36^. The DNA repair response involves activation of ATM through phosphorylation, leading to the activation of several downstream target substrates like γH2A.X and BRCA1. To investigate whether AntiOxCIN4-associated upregulation of cellular antioxidant response impacts the DNA damage response, we examined levels of phosphorylated ATM (p-ATM) and total ATM in treated cells. AntiOxCIN4 treatment led to a robust increase in total and p-ATM, without a corresponding increase in γH2A.X (**Fig. 7h-j**). ATM can phosphorylate BRCA1 and reduce global γH2A.X levels through ubiquitination ^37^. These results prompted us to investigate the BRAC1 gene expression and evidence of H2A.X ubiquitination. In PD fibroblasts treated with AntiOxCIN4, we detected a significant upregulation in BRCA1 levels and increased levels of ubiquitinated H2A.X, which was more abundant than the non-ubiquitinated γH2A.X, thus remaining undetected by Western blot analysis (**Fig. 7k, l)**. Together, these findings demonstrate that AntiOxCIN4 modulates mitochondrial ROS, promotes DNA repair via ATM–BRCA1 signaling, and enhances lysosomal activity, ultimately reducing DNA oxidation and release in EVs. These results mechanistically link mitochondrial ROS, nuclear stress, amphisome-mediated DNA trafficking, and EV-driven inflammation in PD, and identify mitochondrial-targeted antioxidants as a potential therapeutic strategy to modulate EV immunogenicity and inflammation.

## Discussion

Our study identifies a novel pathway linking mitochondrial dysfunction to the extracellular release of oxidized DNA in PD. While mitochondrial ROS and nuclear DNA damage have been separately reported in PD, we provide a mechanistic continuum in which mitochondrial ROS induces γH2AX-positive micronuclei formation, which are subsequently sequestered into amphisomes and exported via small EVs. This positions micronuclei as active intermediates in vesicular cargo loading rather than passive byproducts of genome instability.

Recent work has defined amphisomes as hybrid organelles formed by fusion of multivesicular bodies with autophagosomes, referred to as “amphiectosomes,” and described a common release mechanism for small EVs via limiting membrane rupture ^27^. Extending this concept, we show that oxidatively damaged micronuclei can bypass lysosomal degradation when incorporated into amphisomes, ultimately being expelled as DNA-laden small EVs. This pathway provides an alternative fate for micronuclei in stressed fibroblasts, diverting them away from intracellular cGAS–STING sensing while fueling extracellular inflammatory signaling.

Recent reports indicate that extracellular DNA can be secreted outside of the exosomal pathway via amphisomes ^26,38^. In accordance, we found that oxidized DNA is exported via an alternative route, yielding LC3⁺/CD63⁺ vesicles (amphisome markers), that fall outside the strict definition of classical exosomes. Thus, rather than contradicting earlier studies, this work expands the landscape of vesicular trafficking by uncovering a disease-relevant DNA export mechanism that emerges under mitochondrial and lysosomal stress.

Micronuclei are inherently unstable, and oxidative stress accelerates their collapse and rupture, releasing DNA fragments capable of activating cytosolic cGAS–STING signaling ^18,19^. Our data reveal an alternative fate for oxidatively damaged micronuclei in PD cells, rather than uniformly activating intracellular innate immunity, a fraction are sequestered into amphisomes and expelled through a mechanism common to small EVs. This pathway effectively diverts oxidized nuclear and mitochondrial DNA away from cytosolic sensing, while amplifying extracellular immunogenic signaling. Mitochondrial ROS–induced micronuclear collapse thus emerges as a key upstream driver of vesicular DNA trafficking.

Lysosomal DNA degradation, mediated by LAMP2C, is critical for nuclear and mitochondrial DNA turnover ^39,40^. In PD fibroblasts and postmortem cortical tissue, we observed reduced LAMP2C expression and lysosomal dysfunction, which likely impairs canonical nucleic acid clearance and favors alternative amphisome-mediated routing. This mechanism links mitochondrial ROS and lysosomal stress to enhanced extracellular DNA release, amplifying neuroinflammation in PD. Consistent with prior clinical observations, late-stage PD cerebrospinal fluid exhibits decreased LAMP2 levels correlated with the oxidative stress marker 8-OHdG ^41,42^. Such bypass appears to limit intracellular cGAS activation while enhancing the extracellular release of immunogenic DNA, with oxidized forms being particularly potent at triggering NF-κB–dependent cytokine production in microglia. Thus, the oxidative state of vesicular DNA, rather than its absolute quantity, is a critical determinant of inflammatory potential.

Importantly, the translational relevance of this mechanism is supported by our observation that lysosomal and DNA repair defects identified in PD cortex tissue mirror our cellular observations, including reduced BRCA1 levels and impaired ATM signaling. To our knowledge, this is the first demonstration that in PD micronuclei can evade canonical degradation pathways and be rerouted into amphisomes. This process not only restrains intracellular cGAS activation but also amplifies extracellular immunogenic DNA release, adding a new dimension to EV-mediated neuroinflammation in which the immune activation is not simply triggered by the presence of DNA, but by its biochemical state.

The modulation of mitochondrial redox status using the mitochondrial-targeted antioxidant AntiOxCIN4 ^31^ decreased oxidized DNA loading into small EVs without altering vesicle secretion, restored ATM–BRCA1 DNA repair signaling, and attenuated microglial activation. These results link mitochondrial redox homeostasis to both organelle health and the immunogenic properties of EV cargo, highlighting a novel therapeutic target in PD.

In conclusion, our study uncovers an important new pathway in which mitochondrial ROS and lysosomal dysfunction promote the extracellular export of oxidized DNA–laden small EVs through amphisome-mediated micronuclei sequestration. This mechanism integrates mitochondrial stress, micronuclear collapse, and lysosomal impairment with extracellular immune activation, expanding the functional understanding of EV heterogeneity. By establishing oxidized DNA as a central driver of EV-mediated neuroinflammation and demonstrating its modulation by targeted redox therapy, our study identifies vesicular DNA trafficking as a clinically relevant axis in PD pathogenesis.

### Limitations of the study

A limitation of this study is that our mechanistic analyses were conducted in Parkin-deficient models and sporadic PD patient fibroblasts with mitochondrial dysfunction. Although the sporadic lines lacked known pathogenic genetic alterations, they exhibited elevated mitochondrial ROS production, supporting a role for redox imbalance in altered DNA trafficking. Studies using cells from other genetic forms of PD and a broader panel of idiopathic cases will be necessary to determine the generalizability of this pathway and its relevance to human disease. Future research employing hiPSC-derived neuronal cells and human microglia will be critical to validate whether mitochondrial ROS–driven micronuclei sequestration and amphisome-mediated DNA export represent a shared pathogenic mechanism across PD subtypes. Establishing whether this mechanism is a universal feature of PD or specific to metabolically compromised cases remain an important goal for future work.

## Author contributions

**H.T., O.G., R.G., M.B., CM.D.:** Conducting the experiments, data collection and statistical analysis. **A.B, F.C., F.B., I.M., N.R., P.S., C.K., AR.E., SM.C.:** Provision of materials and samples. **I.M., N.R.:** funding acquisition, review and editing. **P.O., P.P.:** data preprocessing, funding acquisition, quality control and writing—review and editing. **C.L.:** conceptualization, data curation, formal analysis, validation, visualization, funding acquisition, writing—original draft and writing. All authors read and approved the final version of the paper.

## Acknowledgments

We acknowledge Dr Ivan Lalanda and Dr Célia Aveleira for their critical review of the manuscript. Dr. Teresa Rodrigues and Dr. Henrique Girão from the Coimbra Institute for Clinical and Biomedical Research, Faculty of Medicine, University of Coimbra, for the NTA equipment. DNA reporter was kindly provided by Dr Ka Ho Leung (Clarkson University, NY). The authors wish to thank Dr. Margarida Caldeira from the MICC Imaging facility of CNC, partially supported by PPBI – Portuguese Platform of BioImaging (PPBI-POCI-01-0145-FEDER-022122). The authors gratefully acknowledge the IDIBAPS Biobank and Professor I. Ferrer Abizanda from the Bellvitge University Hospital, Institut Català de la Salut, Barcelona, Spain for kindly providing human brain samples and related data to Sandra Morais Cardoso.

## Supplementary Figures

**Figure S1.**
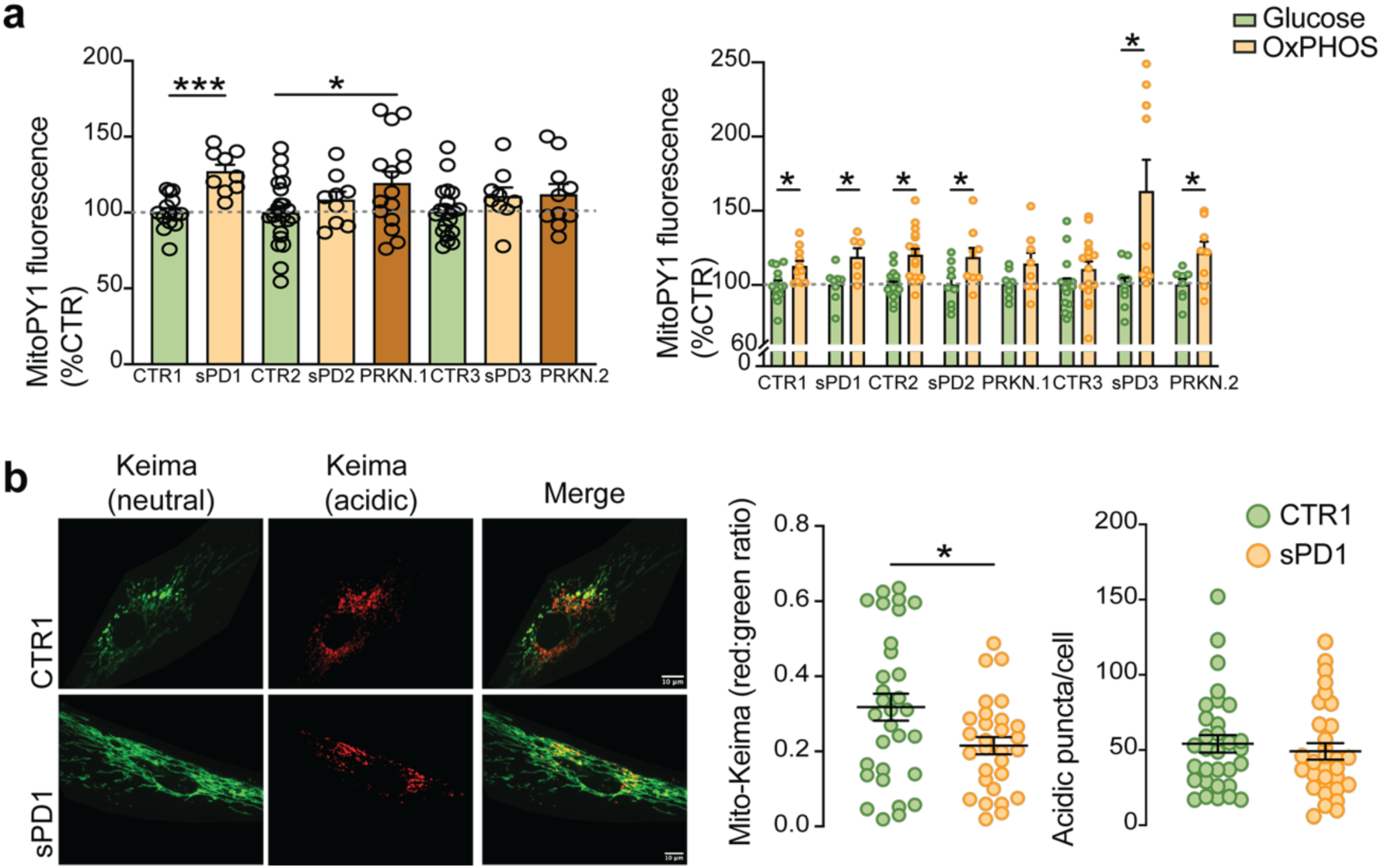
Increased mitochondrial hydrogen peroxide production and impaired mitophagy in fibroblasts from Parkinson’s disease patients. **a,** Mitochondrial hydrogen peroxide levels in MitoPY1-labelled fibroblasts from sporadic PD (sPD) patients and PRKN mutation carriers under basal conditions or forced to rely on oxidative phosphorylation (OXPHOS). **b,** Mitophagy detection with mitochondrial-targeted Keima. Excitation wavelengths of 440 nm (neutral pH) and 590 nm (acidic pH) distinguished mitochondria in the cytosol from those delivered to lysosomes. Representative fluorescence images, red:green fluorescence ratios and counts of acidic Mito-Keima puncta per cell are shown. Scale bar: 10 μm (n = 31). Data are mean ± SEM. Statistical analysis was performed using Welch’s t test (*p < 0.05, ***p ≤ 0.001).

**Figure S2.**
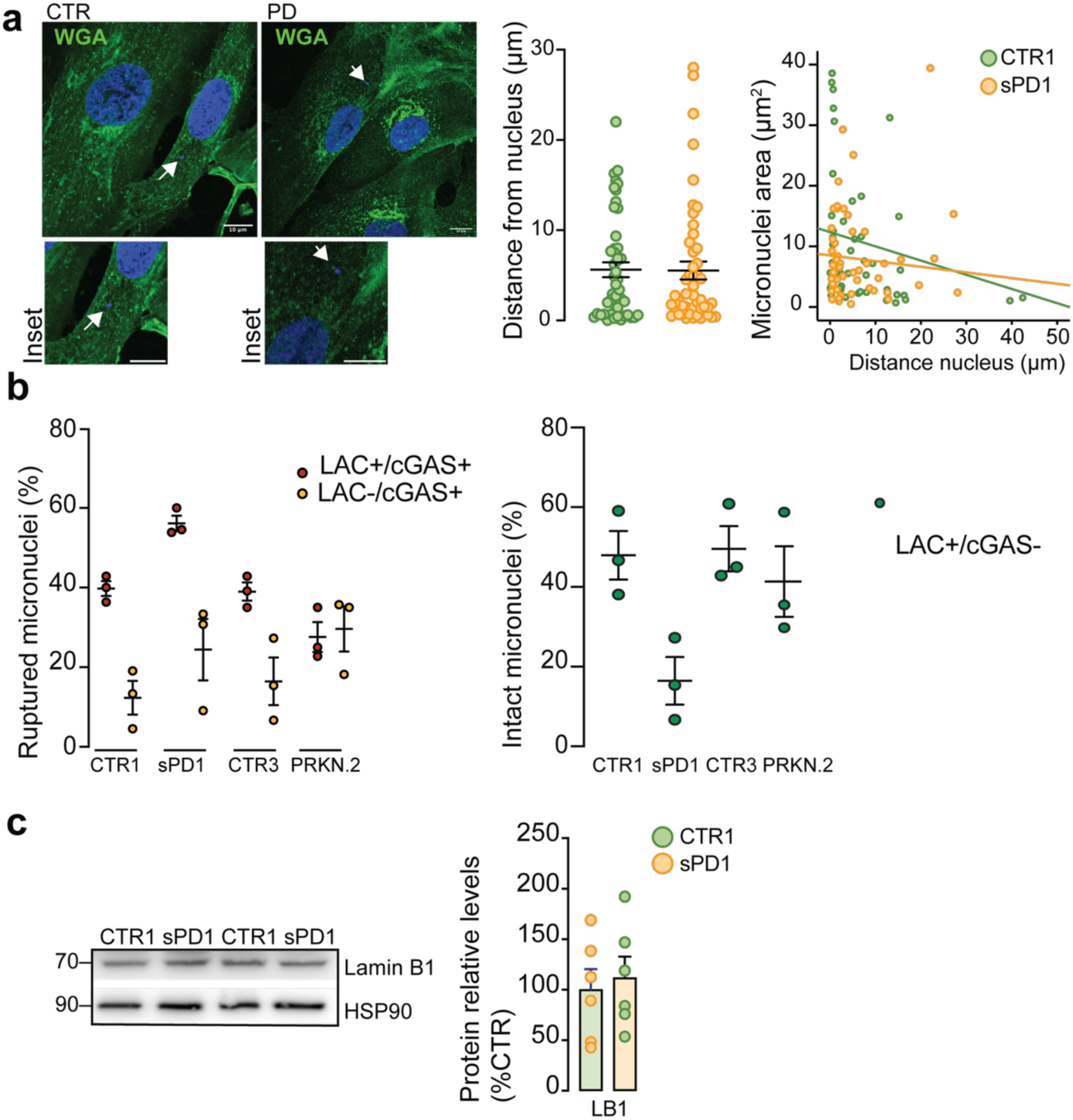
Micronuclei spatial distribution and rupture frequency in PD patient fibroblasts. **a,** Micronucleus–nucleus distance measured using Euclidean distance calculations. Wheat Germ Agglutinin (WGA) staining was used to visualize the cell surface while DAPI was used to label nuclei. Scale bar: 10 μm. A correlation analysis between micronuclear area and distance from the nucleus is shown. **b,** Quantification of ruptured micronuclei (Lamin A/C^+^/cGAS^+^; Lamin A/C^-^) and intact micronuclei (Lamin A/C^+^/cGAS^-^) in fibroblasts from sporadic PD (sPD1) patients and PRKN mutation carriers. **c,** Protein levels of nuclear envelope marker Lamin B1. HSP90 was used as a loading control.

**Figure S3.**
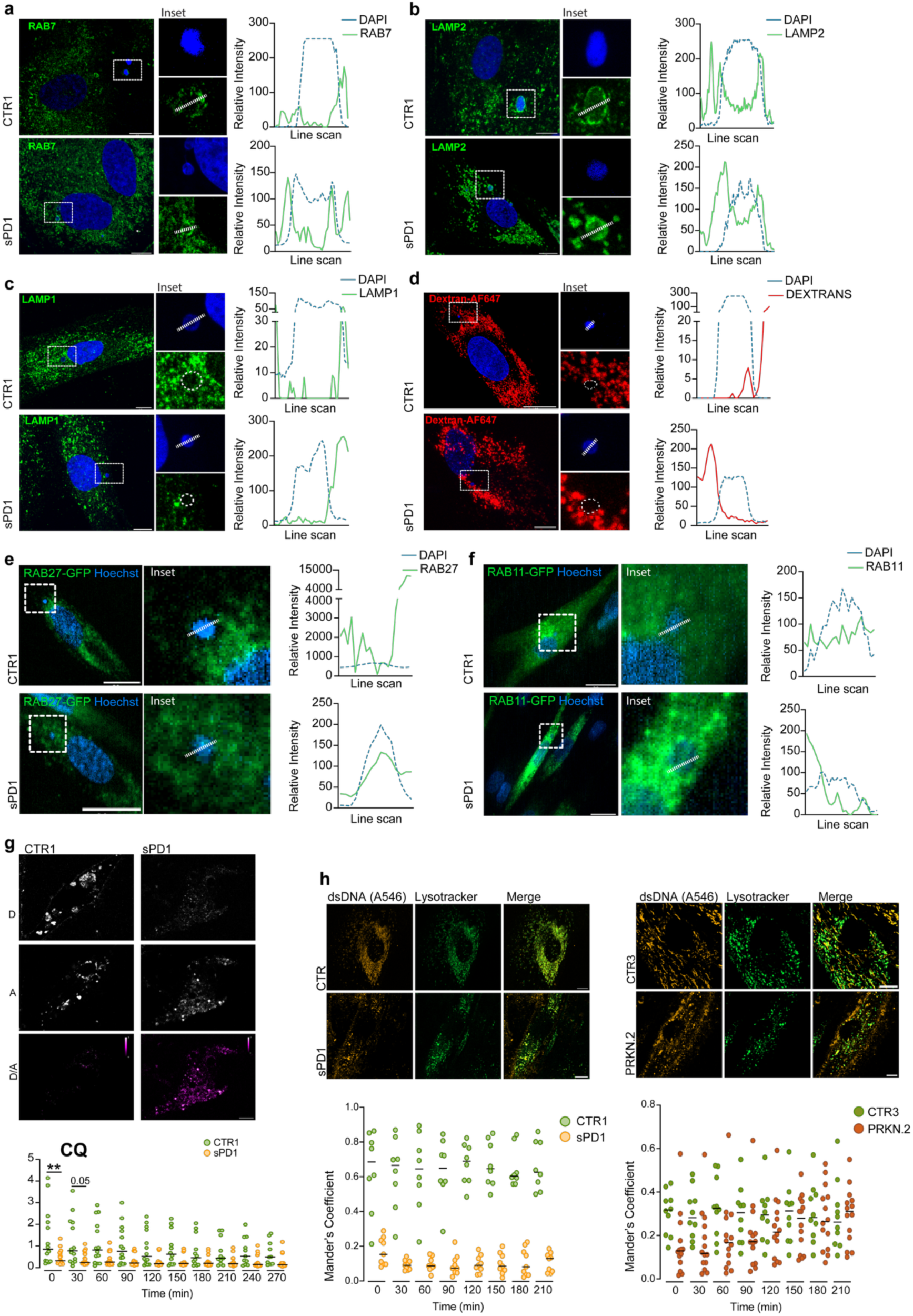
Characterization of endolysosomal markers colocalization with micronuclei. **a–c,** Immunofluorescence of fibroblasts stained for RAB7 (a), LAMP2 (b), and LAMP1 (c), with nuclei counterstained using Hoechst. **d,** Lysosomes labeled by pulse-chase with dextran-AF647; nuclei stained with Hoechst (2 μg/ml). **e-f,** Live-cell imaging of fibroblasts expressing GFP-tagged RAB27 and RAB11. Nuclei were stained with Hoechst (2 μg/ml). Scale: 10 μm and 1μm. Representative images and line-scan profiles. **g,** Quantification of lysosomal D/A ratios over time in cells treated with 100 μM chloroquine (CQ) for 24h. **h,** Quantification of co-localization of DNA reporter with LysoTracker Green (Manders’ Overlap Coefficient: proportion of signal of mono-labeled reporter that is present in Lysotracker channel). Scale: 10 μm. Scatter plot shows individual data points with median. Statistical analysis was performed using unpaired Student’s t-test (*p < 0.05, **p ≤ 0.01).

**Figure S4.**
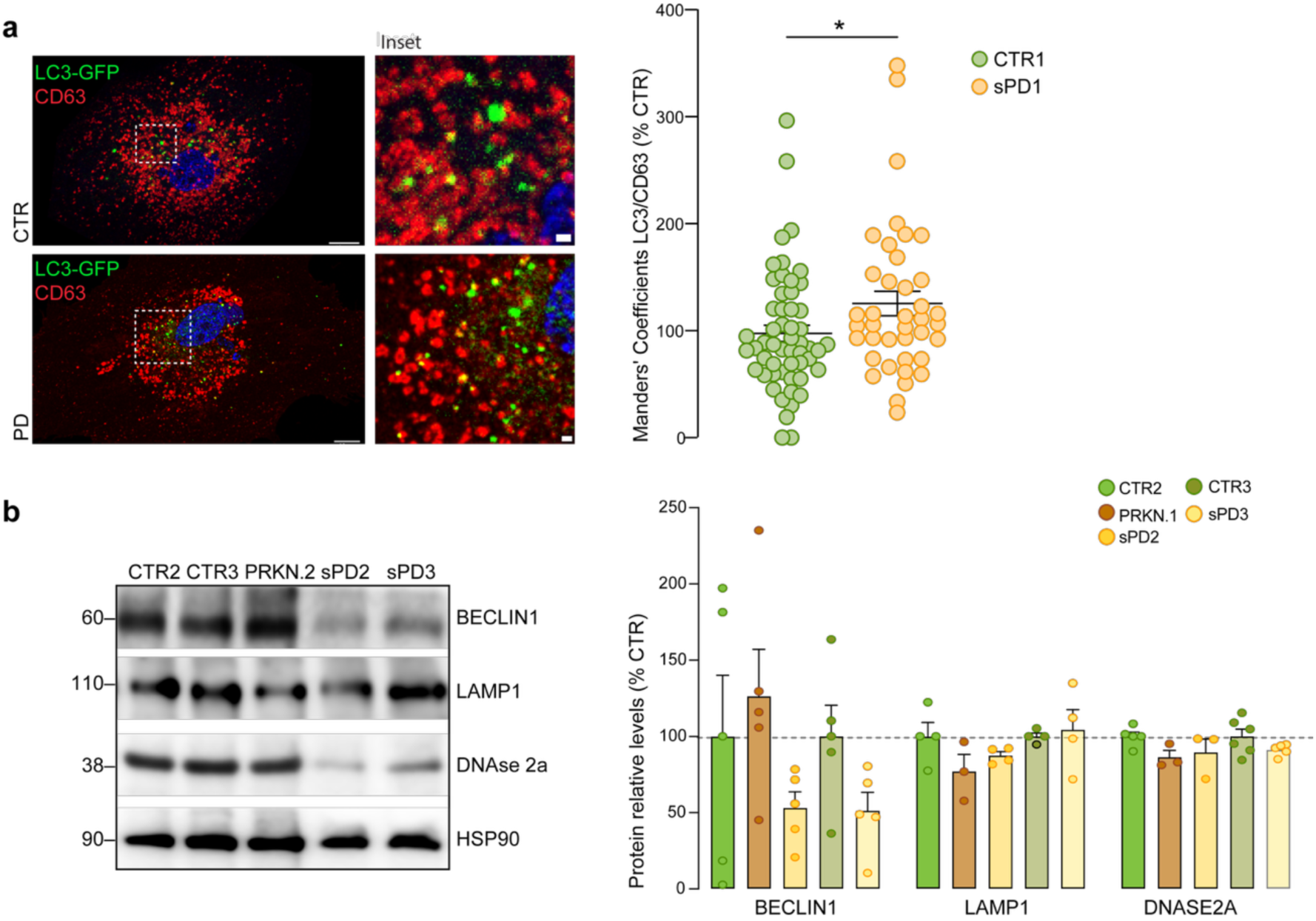
Characterization of autophagic-lysosomal markers. **a,** Colocalization of LC3-GFP⁺ with CD63⁺ in vesicles was quantified using Manders’ coefficient. Representative confocal images and insets are shown (Scale: 10 μm and 1μm). **b**, Protein levels of autophagic-lysosomal markers Beclin1, LAMP1 and DNASE2a. HSP90 was used as loading control. Data are mean ± SEM. Statistical analysis was performed using unpaired Student’s t-test (*p < 0.05).

**Figure S5.**
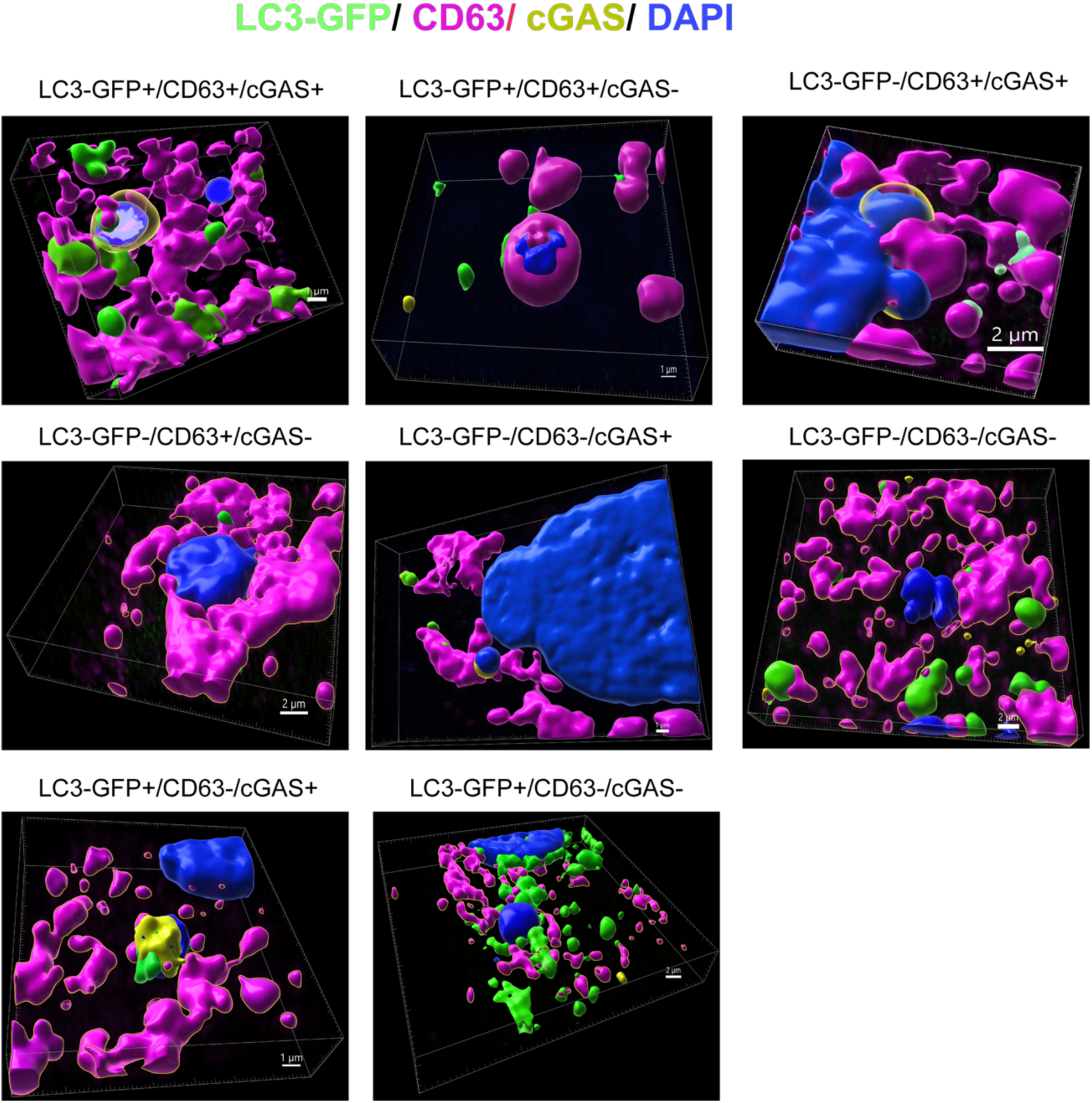
3D rendering of micronuclei categorized by LC3-GFP and CD63 labelling and colocalization with cGAS. Representative Imaris 3D reconstruction illustrating micronuclei classified as LC3-GFP⁺/CD63⁻, LC3-GFP⁻/CD63⁻, LC3-GFP⁻/CD63⁺, and LC3-GFP⁺/CD63⁺, and their colocalization with cGAS.

**Figure S6.**
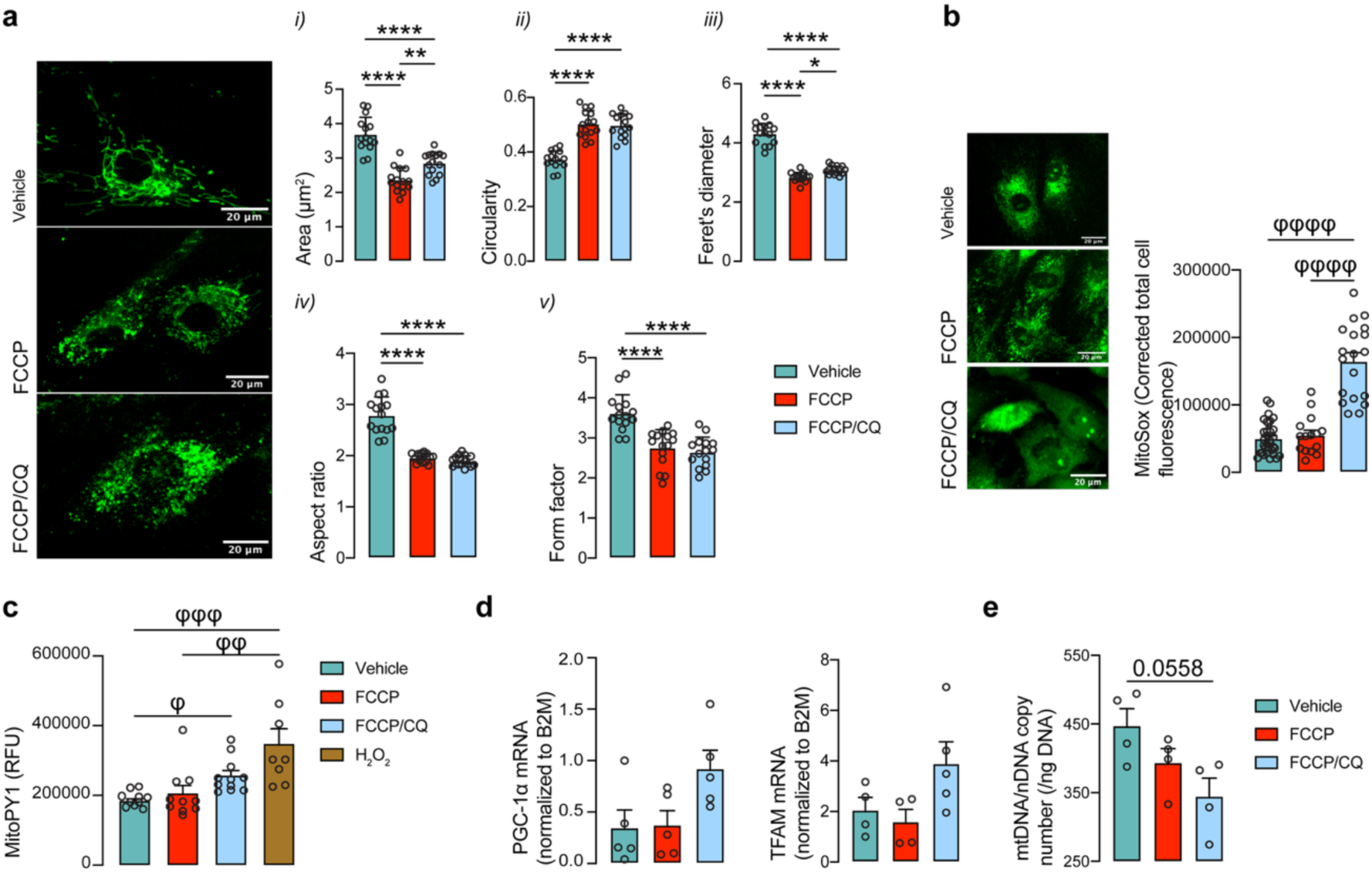
Mitochondria in fibroblasts treated with FCCP and/or CQ to induce dysfunction of the mitochondrial–lysosomal axis exhibit increased fragmentation and elevated production of reactive oxygen species, including superoxide and hydrogen peroxide, along with increased PGC-1α and TFAM expression but reduced mtDNA copy number, indicative of an ineffective compensatory biogenic response. Fibroblasts were treated for 24h with vehicle (DMSO), FCCP (10μM), or CQ (100μM) plus FCCP (10μM). **a,** Quantification of mitochondrial morphometric parameters: i) mitochondrial area per cell normalized to cytoplasmatic area, ii) circularity, iii) Feret’s diameter, iv) aspect ratio and v) form factor and representative images of mitochondria fluorescence after MitoTracker Green staining. Scale bar: 34 μm (n=15). **b,** Detection of mitochondrial superoxide production using MitoSox (corrected total cell fluorescence n=16-35) and of **c,** Hydrogen peroxide production measured in MitoPY1-stained cells (n=26) Representative images of MitoSox-stained cells. Scale bar:52 μm. **d,** mRNA levels of PGC-1α and TFAM normalized to B2M (n=4-5). **e,** mtDNA copy number (n=4). Bar plots represent mean ± SEM. Statistical analysis was performed using one-way ANOVA followed by Tukey’s post hoc test (*p < 0.05, **p < 0.01, ****p < 0.0001) and the Kruskal–Wallis test followed by Dunn’s multiple comparisons test (φ p < 0.05, φφ p < 0.01, φφφ p < 0.001, φφφφ p < 0.0001).

**Figure S7.**
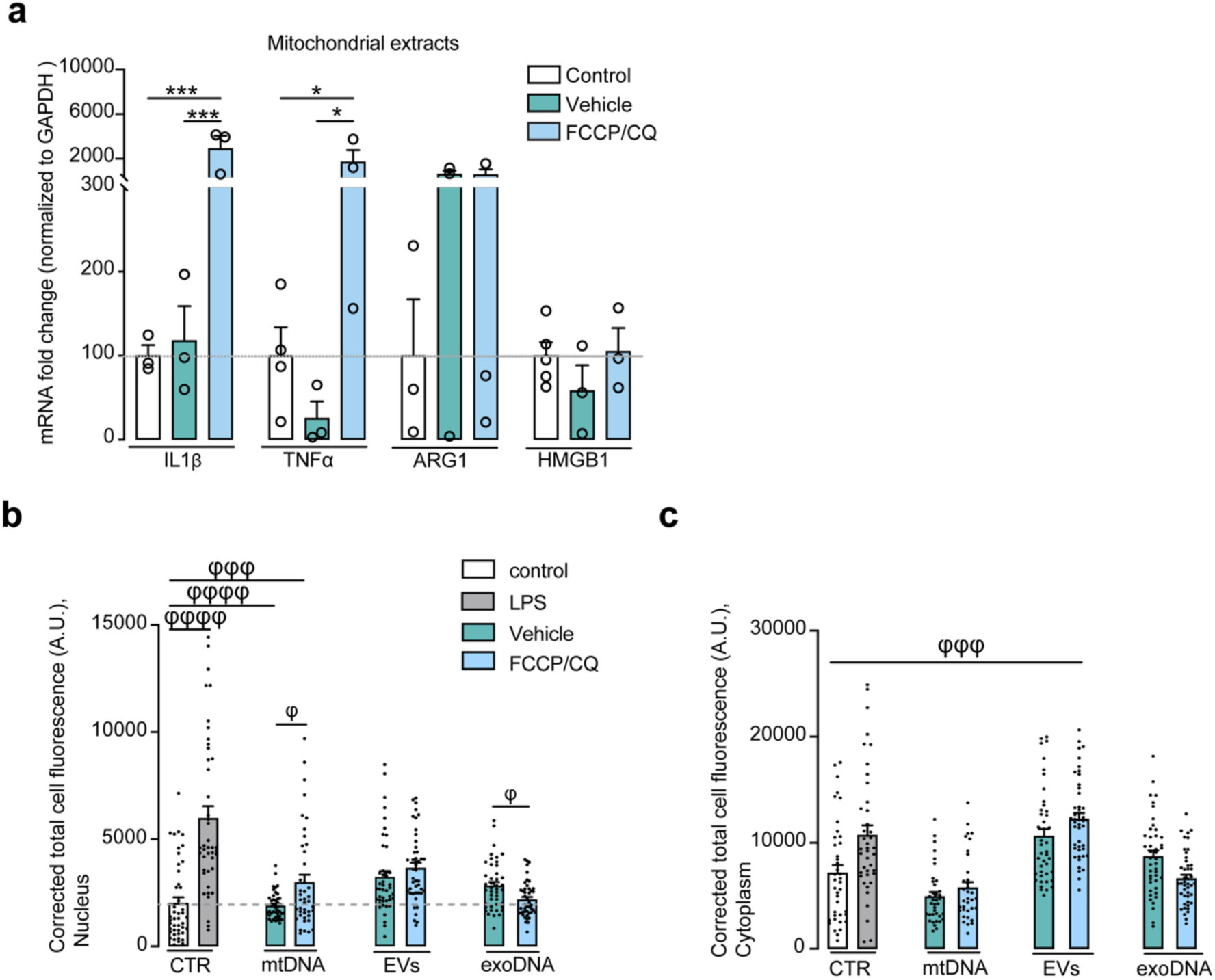
Microglia exposed to intact mitochondria isolated from FCCP/CQ-treated cells exhibited a significantly increased inflammatory response. **a,** Microglia were treated with mitochondria fractions (115 μg) isolated from fibroblasts from each condition. Results are expressed as a percentage of the control value. **b,** Nuclear and cytoplasmatic localization of NF-κB p65 detected by immunofluorescence. Bar plots represent mean ± SEM. Individual data points represent biological replicates. Statistical analysis was performed using a 2-way Anova followed by Tukey’s post hoc test (*p < 0.05, ***p < 0.001), and Kruskal–Wallis test followed by Dunn’s multiple comparisons test (p *φ*< 0.05, *φφφ* p < 0.001, *φφφφ* p < 0.0001).

**Figure S8.**
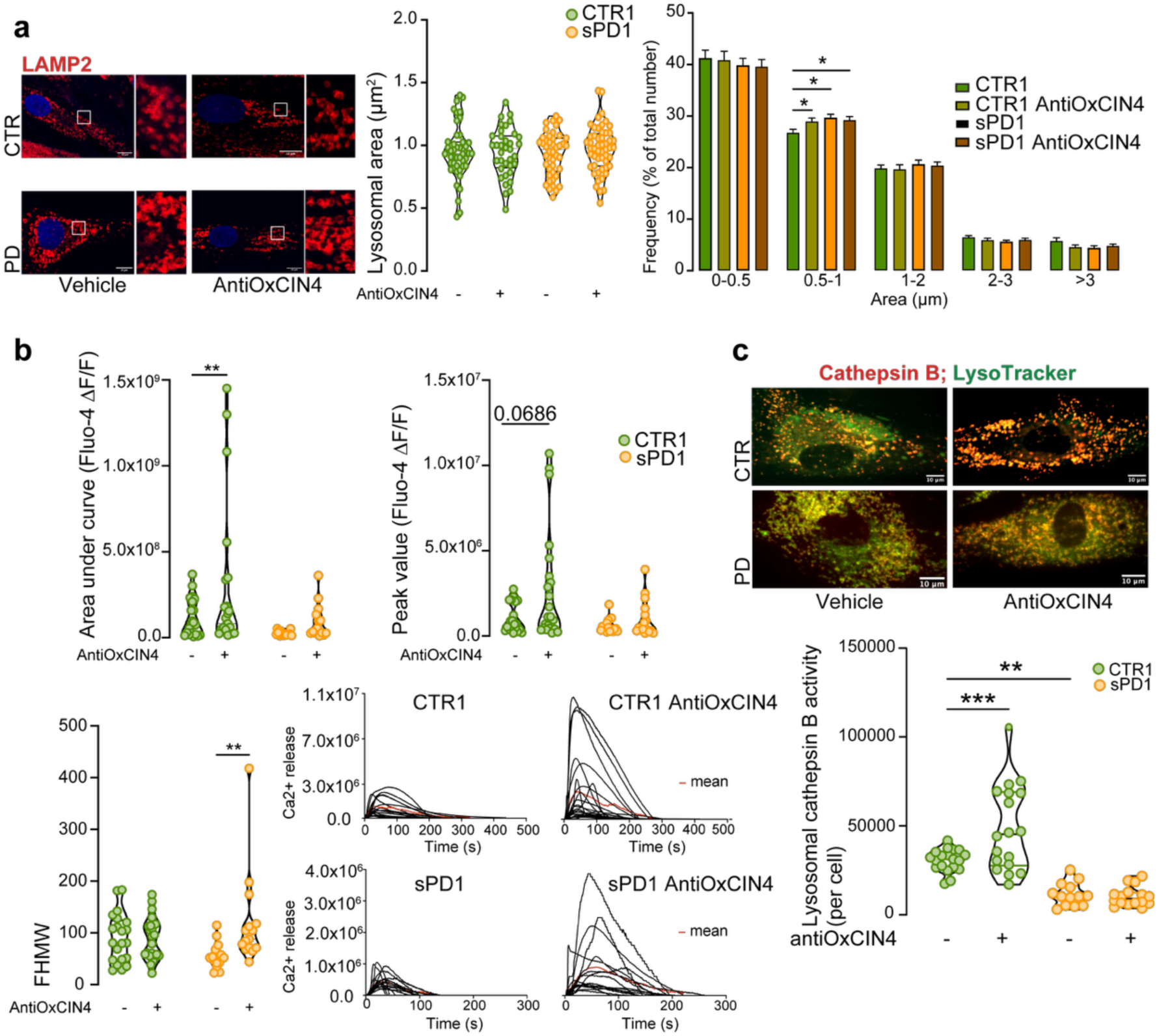
AntiOxCIN4-pretreatment improves lysosomal function. **a,** Representative images of control and PD fibroblasts stained for LAMP2, with quantification of total LAMP2-positive area and histogram of lysosome size distribution. Scale: 10 μm. **b,** Lysosomal Ca²⁺ storage capacity following AntiOxCIN4 treatment. Cells were loaded with FLUO-4 and treated with 400 µM glycyl-L-phenylalanine 2-naphthylamide (GPN) to trigger lysosomal Ca²⁺ release into the cytoplasm. FHMW, full half-maximum width. **c,** Lysosomal proteolytic capacity after AntiOxCIN4 treatment, shown by colocalization of lysosomes with cathepsin B activity in live cells. Scale bar, 10 µm. Individual data points represent biological replicates. Data are mean ± SEM. Statistical analysis was performed using one-way ANOVA followed by Tukey’s multiple comparisons test (*p < 0.05, **p < 0.01, ***p < 0.001).

## References

1 Beatriz, M. et al. Defective mitochondria-lysosomal axis enhances the release of extracellular vesicles containing mitochondrial DNA and proteins in Huntington’s disease. Journal of Extracellular Biology 1, e65 (2022).

2 Knott, A. B., Perkins, G., Schwarzenbacher, R. & Bossy-Wetzel, E. Mitochondrial fragmentation in neurodegeneration. Nat Rev Neurosci 9, 505–518 (2008). 10.1038/nrn2417

3 Xie, Z., Zhang, X., Li, Y. & Zhu, R. Mitochondrial dysfunction drives cellular senescence: Molecular mechanisms of inter-organelle communication. Exp Gerontol 211, 112913 (2025). 10.1016/j.exger.2025.112913

4 Burbulla, L. F. et al. Dopamine oxidation mediates mitochondrial and lysosomal dysfunction in Parkinson’s disease. Science 357, 1255–1261 (2017). 10.1126/science.aam9080

5 Demers-Lamarche, J. et al. Loss of Mitochondrial Function Impairs Lysosomes. J Biol Chem 291, 10263–10276 (2016). 10.1074/jbc.M115.695825

6 Lucien, F. et al. Poly (ADP-Ribose) and α-synuclein extracellular vesicles in patients with Parkinson disease: A possible biomarker of disease severity. PLoS One 17, e0264446 (2022). 10.1371/journal.pone.0264446

7 Peruzzotti-Jametti, L. et al. Neural stem cells traffic functional mitochondria via extracellular vesicles. PLoS Biol 19, e3001166 (2021). 10.1371/journal.pbio.3001166

8 Todkar, K. et al. Selective packaging of mitochondrial proteins into extracellular vesicles prevents the release of mitochondrial DAMPs. Nature communications 12, 1–12 (2021).

9 Hayakawa, K. et al. Transfer of mitochondria from astrocytes to neurons after stroke. Nature 535, 551–555 (2016). 10.1038/nature18928

10 Towers, C. G. et al. Mitochondrial-derived vesicles compensate for loss of LC3-mediated mitophagy. Dev Cell 56, 2029-2042.e2025 (2021). 10.1016/j.devcel.2021.06.003

11 König, T. et al. MIROs and DRP1 drive mitochondrial-derived vesicle biogenesis and promote quality control. Nat Cell Biol 23, 1271–1286 (2021). 10.1038/s41556-021-00798-4

12 McLelland, G. L., Soubannier, V., Chen, C. X., McBride, H. M. & Fon, E. A. Parkin and PINK1 function in a vesicular trafficking pathway regulating mitochondrial quality control. EMBO J 33, 282–295 (2014). 10.1002/embj.201385902

13 Takahashi, A. et al. Exosomes maintain cellular homeostasis by excreting harmful DNA from cells. Nat Commun 8, 15287 (2017). 10.1038/ncomms15287

14 Thakur, B. K. et al. Double-stranded DNA in exosomes: a novel biomarker in cancer detection. Cell Res 24, 766–769 (2014). 10.1038/cr.2014.44

15 Adams, D. J. et al. Genetic determinants of micronucleus formation in vivo. Nature 627, 130–136 (2024). 10.1038/s41586-023-07009-0

16 Welch, G. & Tsai, L. H. Mechanisms of DNA damage-mediated neurotoxicity in neurodegenerative disease. EMBO Rep 23, e54217 (2022). 10.15252/embr.202154217

17 Migliore, L. et al. Oxidative damage and cytogenetic analysis in leukocytes of Parkinson’s disease patients. Neurology 58, 1809–1815 (2002). 10.1212/wnl.58.12.1809

18 Di Bona, M. et al. Micronuclear collapse from oxidative damage. Science 385, eadj8691 (2024). 10.1126/science.adj8691

19 Mackenzie, K. J. et al. cGAS surveillance of micronuclei links genome instability to innate immunity. Nature 548, 461–465 (2017). 10.1038/nature23449

20 Olesen, M. A., Villavicencio-Tejo, F. & Quintanilla, R. A. The use of fibroblasts as a valuable strategy for studying mitochondrial impairment in neurological disorders. Transl Neurodegener 11, 36 (2022). 10.1186/s40035-022-00308-y

21 Qiao, L. et al. LAMP2A, LAMP2B and LAMP2C: similar structures, divergent roles. Autophagy 19, 2837–2852 (2023). 10.1080/15548627.2023.2235196

22 Lan, Y. Y., Londoño, D., Bouley, R., Rooney, M. S. & Hacohen, N. Dnase2a deficiency uncovers lysosomal clearance of damaged nuclear DNA via autophagy. Cell Rep 9, 180–192 (2014). 10.1016/j.celrep.2014.08.074

23 Morse, J. & Leung, K. H. FRET-based reporter assesses lysosomal DNA-degradation ability in live cells. Sensors and Actuators Reports 9 (2025). 10.1016/j.snr.2024.100259.

24 Aizawa, S. et al. Lysosomal membrane protein SIDT2 mediates the direct uptake of DNA by lysosomes. Autophagy 13, 218–222 (2017). 10.1080/15548627.2016.1248019

25 Abu-Remaileh, M. et al. Lysosomal metabolomics reveals V-ATPase- and mTOR-dependent regulation of amino acid efflux from lysosomes. Science 358, 807–813 (2017). 10.1126/science.aan6298

26 Yokoi, A. et al. Mechanisms of nuclear content loading to exosomes. Sci Adv 5, eaax8849 (2019). 10.1126/sciadv.aax8849

27 Visnovitz, T. et al. A ‘torn bag mechanism’ of small extracellular vesicle release via limiting membrane rupture of en bloc released amphisomes (amphiectosomes). Elife 13 (2025). 10.7554/eLife.95828

28 Zhao, M. et al. CGAS is a micronucleophagy receptor for the clearance of micronuclei. Autophagy 17, 3976–3991 (2021). 10.1080/15548627.2021.1899440

29 Xu, Y. & Wan, W. Lysosomal control of the cGAS-STING signaling. Trends Cell Biol 34, 622–625 (2024). 10.1016/j.tcb.2024.05.004

30 Mallach, A. et al. examination of Parkinson’s disease brains suggests decline in mitochondrial biomass, reversed by deep brain stimulation of subthalamic nucleus. FASEB J 33, 6957–6961 (2019). 10.1096/fj.201802628R

31 Deus, C. M. et al. A mitochondria-targeted caffeic acid derivative reverts cellular and mitochondrial defects in human skin fibroblasts from male sporadic Parkinson’s disease patients. Redox Biol 45, 102037 (2021). 10.1016/j.redox.2021.102037

32 Amorim, R. et al. Mitochondriotropic antioxidant based on caffeic acid AntiOxCIN. Free Radic Biol Med 179, 119–132 (2022). 10.1016/j.freeradbiomed.2021.12.304

33 Teixeira, J. et al. Mitochondria-targeted phenolic antioxidants induce ROS-protective pathways in primary human skin fibroblasts. Free Radic Biol Med 163, 314–324 (2021). 10.1016/j.freeradbiomed.2020.12.023

34 Morgenstern, C. et al. Biomarkers of NRF2 signalling: Current status and future challenges. Redox Biol 72, 103134 (2024). 10.1016/j.redox.2024.103134

35 Ong, A. J. S. et al. The KEAP1-NRF2 pathway regulates TFEB/TFE3-dependent lysosomal biogenesis. Proc Natl Acad Sci U S A 120, e2217425120 (2023). 10.1073/pnas.2217425120

36 Lee, J. H. et al. ATM directs DNA damage responses and proteostasis via genetically separable pathways. Sci Signal 11 (2018). 10.1126/scisignal.aan5598

37 Krum, S. A., la Rosa Dalugdugan, E., Miranda-Carboni, G. A. & Lane, T. F. BRCA1 Forms a Functional Complex with γ-H2AX as a Late Response to Genotoxic Stress. J Nucleic Acids 2010 (2010). 10.4061/2010/801594

38 Jeppesen, D. K. et al. Reassessment of Exosome Composition. Cell 177, 428–445.e418 (2019). 10.1016/j.cell.2019.02.029

39 Fujiwara, Y., Hase, K., Wada, K. & Kabuta, T. An RNautophagy/DNautophagy receptor, LAMP2C, possesses an arginine-rich motif that mediates RNA/DNA-binding. Biochem Biophys Res Commun 460, 281–286 (2015). 10.1016/j.bbrc.2015.03.025

40 Hase, K. et al. RNautophagy/DNautophagy possesses selectivity for RNA/DNA substrates. Nucleic Acids Res 43, 6439–6449 (2015). 10.1093/nar/gkv579

41 Klaver, A. C., Coffey, M. P., Aasly, J. O. & Loeffler, D. A. CSF lamp2 concentrations are decreased in female Parkinson’s disease patients with LRRK2 mutations. Brain Res 1683, 12–16 (2018). 10.1016/j.brainres.2018.01.016

42 Boman, A. et al. Distinct Lysosomal Network Protein Profiles in Parkinsonian Syndrome Cerebrospinal Fluid. J Parkinsons Dis 6, 307–315 (2016). 10.3233/JPD-150759

